# Scalable identification of lineage-specific gene regulatory networks from metacells with NetID

**DOI:** 10.1101/2024.09.08.611796

**Authors:** Weixu Wang, Yichen Wang, Ruiqi Lyu, Dominic Grün

## Abstract

The identification of gene regulatory networks (GRN) governing distinct cell fates in multilineage cellular differentiation systems is of critical importance for understanding cell fate decision. Single-cell RNA-sequencing (scRNA-seq) provides a powerful tool for the quantification of gene-level co-variation across the cell state manifold. However, accurate GRN reconstruction is hampered by the sparsity of scRNA-seq data introducing substantial technical noise. Moreover, the high dimensionality of typical scRNA-seq datasets limits the scalability of available approaches. To overcome these challenges, and to facilitate the inference of lineage-specific GRNs with directed regulator-target relations, we introduce NetID. This approach optimizes coverage of the cell state manifold by homogenous metacells and avoids spurious gene-gene correlations observed with available imputation methods. Benchmarking demonstrates superior performance of NetID compared to imputation-based GRN inference. By incorporating cell fate probability information, NetID facilitates prediction of lineage-specific GRNs and recovers known network motifs centered around lineage-determining transcription factors governing bone marrow hematopoiesis, making it a powerful toolkit for deciphering the gene regulatory control of cellular differentiation from large-scale single-cell transcriptome data.

## Introduction

The development of large-scale scRNA-seq techniques over the past decade has revolutionized our ability to unbiasedly discriminate cell states based on the transcriptome fingerprint of individual cells. Numerous computational methods facilitate the inference of cellular differentiation trajectories from snapshot or time-course scRNA-seq datasets [1]. Such approaches typically rely on differentiation pseudotime estimation and permit the analysis of gene expression dynamics underpinning cell fate decision and terminal differentiation.

Exploiting covariation of the expression patterns of individual genes permits the inference of gene regulatory networks (GRNs) encoding regulator-target relations in a system of interest. Multiple approaches for systematic GRN inference have been introduced in the past. One of the most widely used methods, GENIE3 [2], utilizes random forest regression for GRN construction and has shown favorable performance in a recent benchmarking study [3], while another top-performing method, PIDC [4], relies on partial information decomposition.

Although these methods successfully recover known regulatory interactions, sensitivity and specificity of inferred regulatory links suffer from a high level of technical noise due to the sparsity of scRNA-seq data, which has been identified as one of the major challenges in the analysis of such data [5].

To overcome the problem of sparsity, or dropout, i.e., the absence of gene read counts as a consequence of prevalent sampling noise, various computational methods have been proposed for imputing missing readout from the gene expression information of each cell’s neighborhood in order to smoothen inferred gene expression across the cell state manifold [6-9]. However, these approaches typically induce spurious correlations between the expression levels of different genes, leading to a decrease in GRN reconstruction performance [10-12].

Another solution to alleviate technical noise caused by data sparsity relies on the utilization of metacells [13]. The concept of metacells relies on local groups of cells considered to be sampled from the same state, covering the cell state manifold. This approach was proposed as a way of maintaining statistical utility while maximizing effective data resolution. Although metacells are far more granular than clusters and are optimized for homogeneity, the metacell inference on unpruned cell k-nearest neighbor (KNN) graphs without explicitly testing for cell state homogeneity based on a background model capturing known noise components [14, 15] may lead to a mixing of distinct cell states within individual metacells and therefore compromise GRN inference.

Finally, available methods for GRN inference do not account for differences in GRN architecture across distinct lineages within multilineage scRNA-seq data.

To overcome the challenge of technical noise and to facilitate accurate and scalable inference of lineage-specific GRNs, we introduce NetID. The NetID algorithm builds on the metacell concept applied to pruned KNN graphs. We demonstrate that NetID preserves biological covariation of gene expression, and outperforms GRN inference with imputation-based methods. By incorporating cell fate probability information, we enable the inference of cell-lineage specific GRNs, which permit the recovery of ground truths network motifs driven by known lineage-determining transcription factors of mouse hematopoietic bone marrow cells.

NetID provides a novel toolkit to infer gene regulatory network in large-scale single-cell gene expression data.

## Results

### NetID algorithm for scalable inference of lineage-specific GRNs

To circumvent the problem of data sparsity due to sampling dropouts of sequenced mRNAs in individual cells, the concept of metacells has been introduced [13]. Metacells are defined as disjoint homogenous groups of cells sampled from the same distribution. NetID provides a novel GRN inference method relying on metacells in order to (1) facilitate scalability of GRN inference to large single-cell datasets, and (2) limit the adverse effect of data sparsity on the inference of gene-gene covariation underlying GRNs. In order to identify a limited number of metacells capturing all relevant variability defining the cell state manifold of a given scRNA-seq dataset, NetID first performs sampling of cells after normalization and transformation by principal component analysis (PCA) utilizing a sketch-based method called geosketch with the objective to obtain homogenous coverage [16].

These sampled cells are defined as “seed cells”. For the inference of metacells, NetID starts by computing the k-nearest neighbors (KNN) of each seed cell. To only keep cells consistent with sampling from the same distribution for each metacell, outlier cells are pruned from each neighborhood based on a local background model of gene expression variability as implemented in VarID2 [15]. VarID2 computes the probability of observing a gene-specific transcript count for each of the neighbors according to a negative binomial distribution parametrized by the local mean-variance dependence across all genes. If the expression in a nearest neighbor cell is significantly different from the seed cell, VarID2 assigns a low probability to the corresponding edge to enable pruning of the KNN graph by applying a p-value cutoff. This procedure removes unwanted variability arising from the admixture of distinct cell states and maximizes homogeneity of metacells.

To avoid inflation of gene-gene covariance as a result of overlapping metacells and to ensure that the inferred metacells represent independent states on the manifold we avoid occurrence of the same cell in the neighborhoods of different seed cells.

To achieve this, we assign a shared neighbor to the seed cell with the largest edge p-value. Shared neighbors remaining after this step are assigned to the seed cell with the lowest number of neighbors. Remaining neighbors are termed partner cells. Metacells with too few partner cells are removed. To obtain the expression profile of each metacell, normalized or raw gene counts are aggregated (Fig .1).

**Figure 1.**
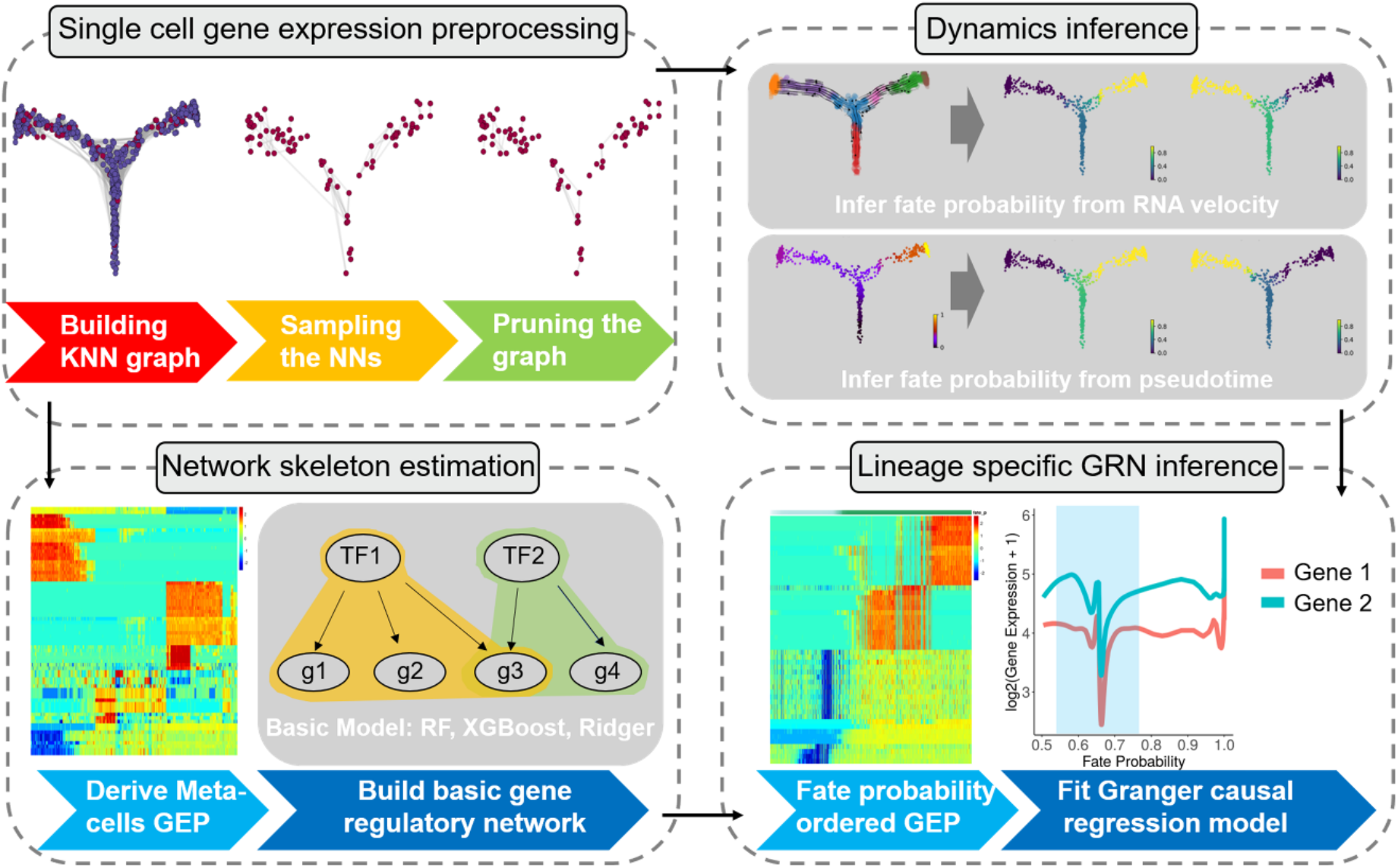
The NetID algorithm. NetID utilizes metacells to infer gene regulatory networks (GRNs) from large single-cell datasets to increase scalability and reduce technical noise. First, NetID performs cell sampling using geosketch and applies VarID2 to prune the KNN graph. Subsequently, cells are ordered according to cell fate probability and regulator-target interactions are inferred by Granger ridge regression. By integrating the GRN inferred from the Granger causal model and GENIE3, NetID learns lineage-specific GRNs to identify key regulators of cell fate decision. KNN: k-nearest neighbor; NN: nearest neighbor; GEP: gene expression profile. RF: random forests.

The NetID approach is designed to infer metacells capturing predominant cell state variation across the manifold, while reducing the sample size needed for accurate gene regulatory network (GRN) inference. NetID integrates GENIE3 for GRN inference, but alternative methods can be applied on metacell profiles. However, since each cell lineage is potentially governed by a unique GRN topology, a global GRN model may provide insufficient resolution or even confound lineage-specific sub-networks. To overcome this limitation, we utilize cell fate probability inferred from pseudotime [17] or RNA velocity [18, 19] to order cells along their respective lineage trajectories. This allows the prediction of directed regulator-target gene relations by ridge regression Granger causality tests [20]. By integrating the GRN inferred from the Granger causal model and GENIE3, we can learn lineage-specific GRNs that enable identification of important driver genes and regulatory interactions during cell fate decisions.

### Testing the NetID architecture on a ground truths dataset

To investigate the contribution of each step in the NetID algorithm towards GRN prediction performance, we conducted testing on a hematopoietic differentiation cell dataset [21] using a GRN curated from nonspecific ChIP-seq data [3] as ground truth. Comparing random (Fig. 2A) and geosketch sampling (Fig. 2B) of seed cells demonstrated that geosketch sampling led to smaller Hausdorff distance to all other cells (Fig. 2C) and explained more gene expression variation (Fig. 2D), consistent with previous findings [16]. We confirmed these observations on human adult hematopoietic differentiation [22] and embryonic stem cell [23] datasets (Figure S2). Then compared the two sampling methods by directly aggregating k-nearest neighborhoods of each seed cell without pruning and reassignment prior to GRN inference by GENIE3. Early precision rate (EPR) and area-under-receiver-operating-characteristic curve (AUROC) metrics indicated significantly improved GRN inference of geosketch compared to random sampling (Fig. 2E left panel, Fig. S3A).

**Figure 2.**
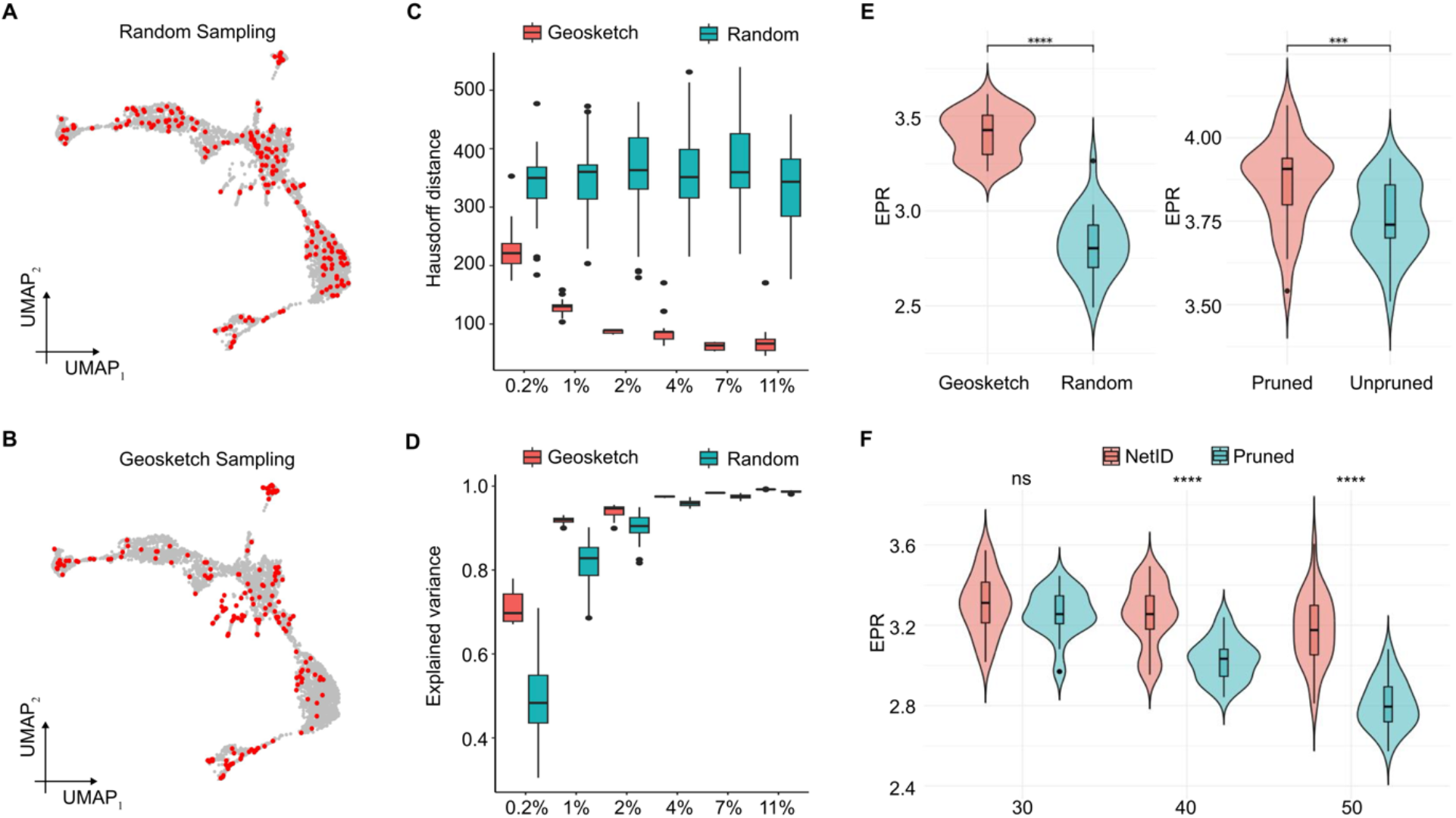
Validating GRN inference performance of NetID on hematopoietic ground truths data. A) UMAP representation of Kit+ hematopoietic progenitors [21] highlighting randomly sampled cells in red. B) UMAP representation of Kit+ hematopoietic progenitors [21] highlighting geosketch sampled cells in red. C) Boxplot comparing the Hausdorff distance of randomly sampled cells (green) and geosketch sampled cells (red) with 30 repeats. The x-axis denotes the fraction of sampled cells and the absolute number in parentheses. D) Boxplot comparing the explained expression variance (Methods) of randomly sampled cells (green) and geosketch sampled cells (red) with 30 repeats. The x-axis denotes the fraction of sampled cells and the absolute number in parentheses. E) Violinplots showing the difference in early precision rate when using GENIE3 inferred GRN on geosketch sampled cells or randomly sampled cells (left panel) with 30 repeats, and when using GENIE3 inferred GRN on pruned or unpruned KNN graphs (right panel). F) Violinplots showing the comparison of early precision rate between two different strategies. “NetID” denotes combining geosketch, KNN graph pruning, and neighbor reassignments to build metacell profiles for GRN inference with 30 repeats. “Pruned” denotes combining geosketch and KNN graph pruning to build metacell profiles for GRN inference with 30 repeats. The x-axis denotes the number of neighbors. We benchmarked the performance on the non-specific ChIP-seq network as the ground truth. In (E-F), the box in the violinplot represents the interquartile range (IQR). The whiskers extend to the smallest and largest values within 1.5 times the IQR. The black line within the box indicates the median.. *P<0.05, **P<0.01, ***P<0.001, two-sided Wilcoxon rank sum test.

To demonstrate the benefit of neighbor pruning, we performed direct cell aggregation on pruned and unpruned KNN graphs without additional reassignment of shared partner cells for geosketch-sampled seed cells, and observed significantly improved GRN inference performance when pruning was included (Fig. 2E right panel, Fig. S3A). We repeated these analyses using the STRING database to derive a ground truth network, and could confirm the observed improvement (Fig. S3B,C).

Finally, we compared cell aggregation on pruned KNN graphs with and without reassignment of shared partner cells, and benchmarked the performance at varying neighborhood sizes. Together with the previous tests, this comparison indicates that NetID, combining geosketch sampling, KNN graph pruning and shared partner cell reassignment, improves GRN inference performance. These observations were consistent across datasets as demonstrated by extensive step-by-step benchmark on the human hematopoietic differentiation and embryonic stem cell datasets (Fig. S4-6). For the latter, we included an available cell specific ChIP-seq network (Methods) from BEELINE [3] as ground truth.

A critical parameter of NetID is the optimal number of seed cells. Considering the entire metacell inference process, we observed that increasing the seed cell sample size decreases the number of partner cells for each seed cell (Fig. S7). The required number of sampled seed cells depends on the cell state heterogeneity of the dataset, while the number of partner cells controls the sparsity of the inferred metacell profiles. Optimal GRN inference from metacell profiles relies on sufficient coverage of the cell state manifold by seed cells with a limited degree of metacell sparsity to avoid sampling noise. Combining both objectives into a score (Methods) enables the inference of a “sweet spot” for the optimal number of sampled seed cells. We evaluated the GRN inference accuracy with varying numbers of seed cells and confirmed that the predicted “sweet spot” matched well with the performance optimum in terms of EPR and AUROC (Fig. S8).

We further observed that removal of seed cells with a low number of partner cells after pruning and reassignment improves GRN inference (Fig. S9), suggesting that careful inference of metacell gene expression improves the recovery of gene-gene covariation.

### Benchmark of NetID with available imputation methods

To benchmark the influence of NetID’s metecall generation step against other inference strategies with imputation-based methods including DCN [6], MAGIC [8], SAVER [7], KNN (mean expression across KNN), or with raw transcript counts based on *in silico* generated ground truths data, we used dyngen [24] to perform scRNA-seq data simulation. This allowed us to determine the ground truth GRN. We simulated two scRNA-seq datasets with different topological structures, i.e., bifurcating (Fig. 3A) and cyclical (Fig. 3C) topology. We then used EPR, AUPRC, and AUROC to evaluate the performance of GRN inference. Overall, NetID exhibited superior performance reflected by all metrics and was always among the top methods when compared to imputation-based approaches (Fig. 3B, D). Overall, NetID provided competitive GRN inference performance on simulated datasets and was more robust compared to raw gene expression data and imputation-based methods.

**Figure 3.**
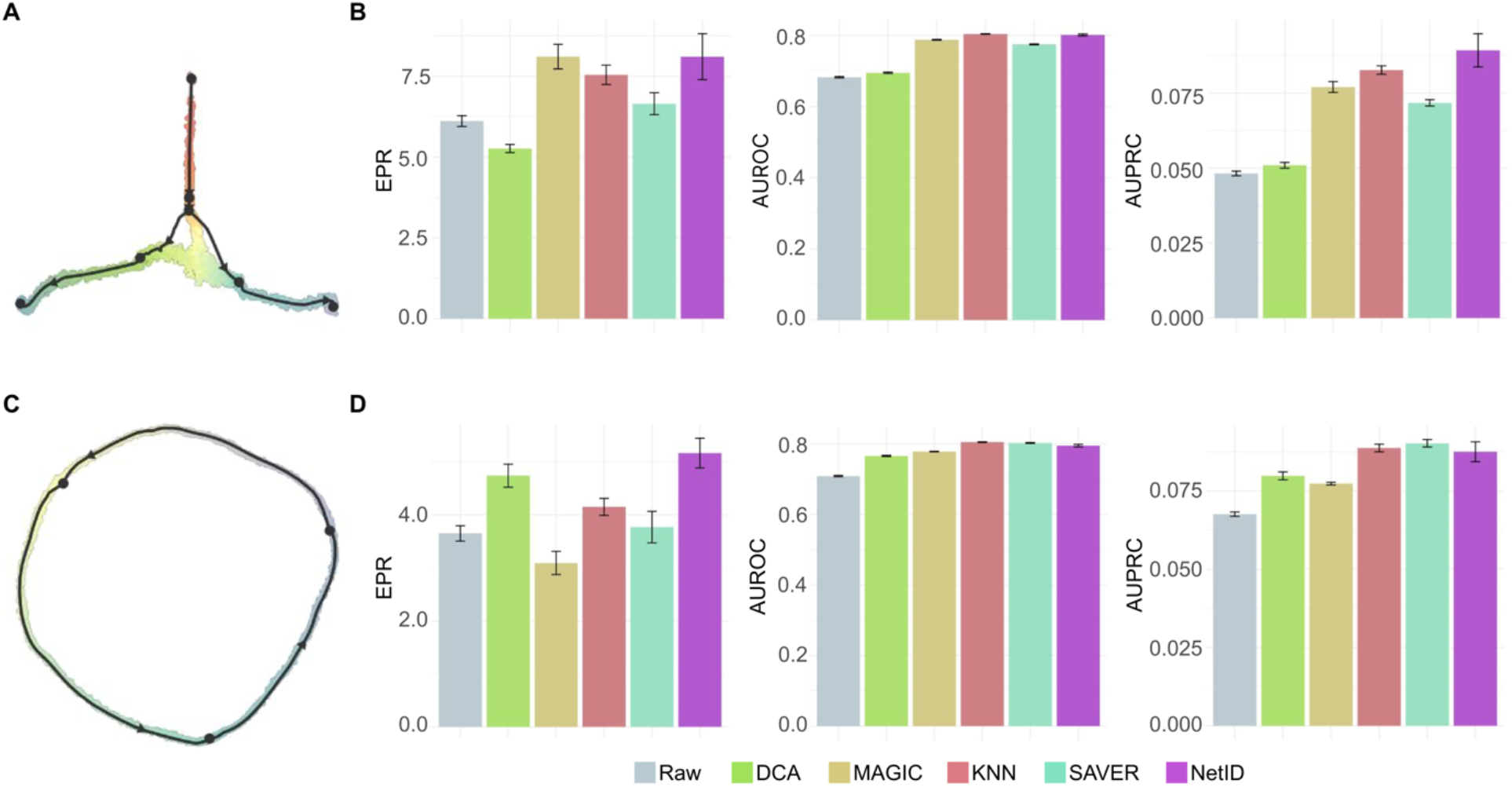
Benchmarking of NetID on simulated datasets. A) Dimensional reduction representation of a simulated scRNA-seq dataset with bifurcating trajectories. B) Barplot showing the early precision rate (EPR) (left panel), area under the receiver operating characteristic curve (AUROC) (middle panel), and area under the precision-recall curve (AUPRC) (right panel) of the GRN inferred from raw gene expression profiles, imputed gene expression profiles using four methods (DCA, MAGIC, KNN and SAVER), and metacell expression profiles inferred by NetID for data in (A). NetID was run 30 times to generate error bars. For other imputation methods, we randomly sampled 90% of cells each time to infer performace error bars. GENIE3 was used for GRN inference. C) Dimensional reduction representation of a simulated scRNA-seq dataset with a cycling trajectory. D) Barplot showing the EPR (left panel), AUROC (middle panel), and AUPRC (right panel) of the GRN inferred from raw gene expression profiles, imputed gene expression profiles using four methods (DCA, MAGIC, KNN and SAVER), and metacell expression profiles inferred by NetID for data in (C). NetID was run 30 times to generate error bars. GENIE3 was used for GRN inference. For other imputation methods, we randomly sampled 90% of cells each time to infer performace error bars.

Since simulation-based methods do not fully reflect the complexity of real scRNA-seq data, we further utilized the BEELINE GRN benchmarking pipeline [3] to compare NetID with the same imputation-based methods and *MetaCell* [13] based inference on real scRNA-seq datasets for mouse embryonic stem cell (mESC) differentiation [23] and mouse hematopoietic stem cell (mHSC) differentiation[25]. For both systems, we selected ground truths networks inferred from ChiP-seq data specific to the respective system or from non-specific ChIP-seq or STRING datasets following the strategy of BEELINE [3]. In case of mESCs, we further utilized ground truths based on loss-of-function/gain-of-function data [3].Overall, NetID exhibited superior performance compared to raw gene expression and imputation-based methods for both systems according to EPR values (Fig. 4A-B). The only exception was the benchmarking of HSC differentiation based on HSC-specific ChiP-seq data. However, none of the methods outperformed random guessing (EPR=1) in this case, suggesting that this ground truth network may not fit the real underlying network in the tested mHSC datasets.

**Figure 4.**
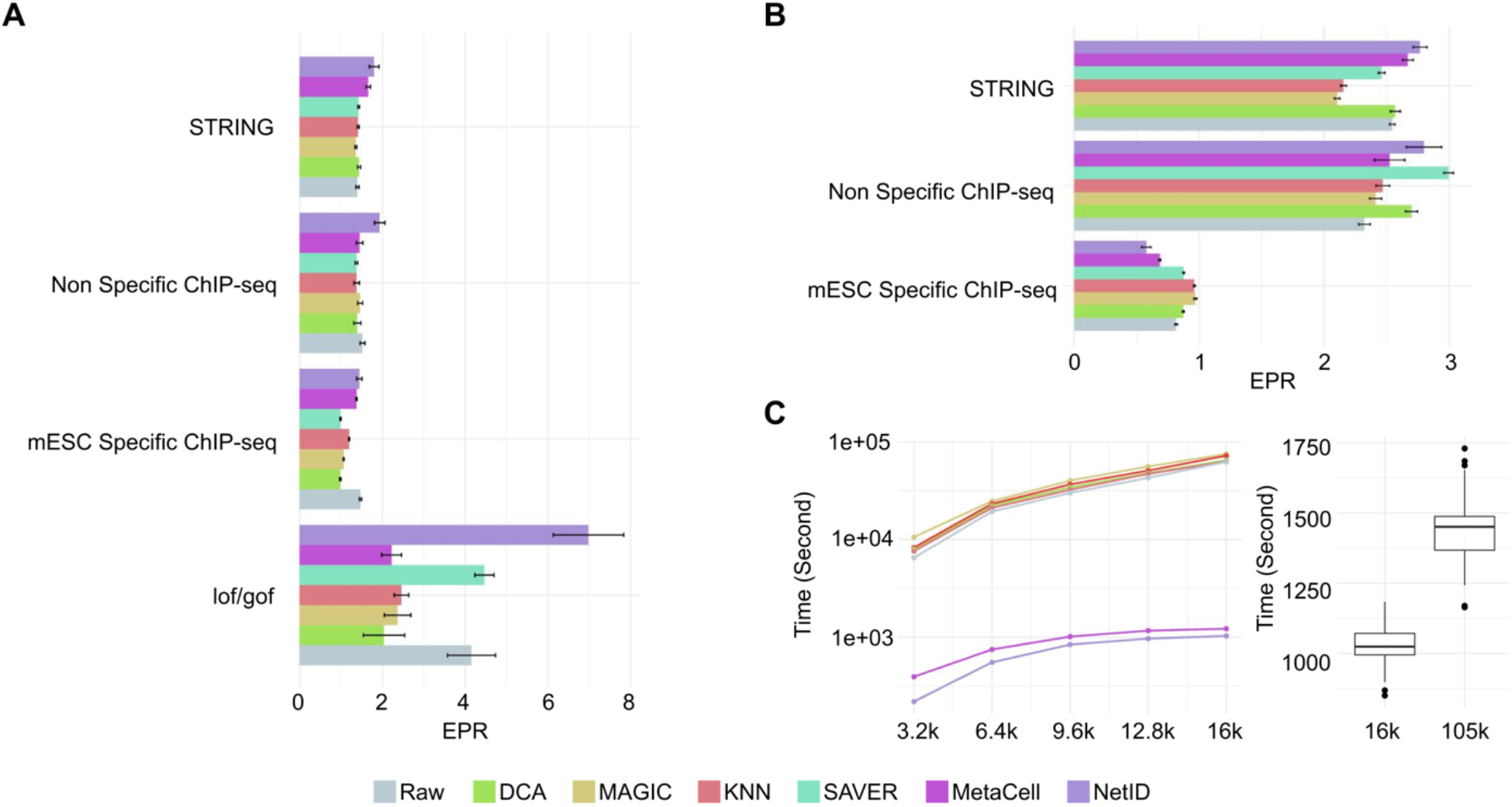
Benchmarking gene regulatory network inference on real scRNA-seq dataset. A) Barplot showing the early precision rate of the gene regulatory network (GRN) inferred from raw gene expression, imputed gene expression using four alternative methods, and metacell gene expression profiles (MetaCell and NetID) for mouse embryonic stem cell data [23]. NetID was run 30 times to generate error bars. For other imputation methods, we randomly sampled 90% of cells each time to infer performace error bars. Lof/gof: loss-of-function/gain-of-function B) Barplot showing the early precision rate of the GRN inferred from raw gene expression, imputed gene expression using four alternative methods, and metacell gene expression profiles (MetaCell and NetID) for mouse hematopoietic stem cell data [25]. NetID was run 30 times to generate error bars. For other imputation methods, we randomly sampled 90% of cells each time to infer performace error bars. C) Line plot showing the running time of the GENIE3 algorithm on raw gene expression profiles and processed gene expression generated from six methods for a subset of 16k cells extracted from mouse embryogenesis datav[26] with different percentages of the cells (left panel). The boxplot displays the running time comparison when NetID was applied to a larger subset with 105k cells compared to the 16k cell dataset with 30 repeats. The benchmarking was performed on a workstation with 16 Intel(R) Xeon(R) Gold 6242 CPUs and 128 GB RAM. In (C), the box in the boxplot represents the interquartile range (IQR). The whiskers extend to the smallest and largest values within 1.5 times the IQR. The black line within the box indicates the median.. *P<0.05, **P<0.01, ***P<0.001, two-sided Wilcoxon rank sum test.

A critical bottleneck for GRN inference is computational speed. We compared the computation speed of the different imputation methods and calculated the running time as the sum of each algorithm’s running time and the running time of GENIE3 network inference on the large scale mouse embryogenesis atlas dataset [26]. As expected, metacell-based methods, i.e., NetID and MetaCell, were the fastest among all methods. In contrast, the imputation-based methods required a long time to perform both imputation and GRN construction (Fig. 4C, left). Specifically, NetID was 10 times faster than any other imputation-based method. While GENIE3 cannot handle large datasets with ∼100k, we show that application of NetID to a ∼105k mouse embryogenesis dataset requires <25min running time (Fig. 4C, right).

Overall, NetID’s strategy of generating metacell gene expression profiles facilitates superior GRN inference performance compared to raw gene expression profiles and imputation-based methods according to the benchmark results on simulated and real datasets. In particular, due to its fast computation speed NetID enables scalability of GRN inference to large-scale scRNA-seq datasets.

### Incorporating cell fate probability for lineage-specific GRN inference

In a multilineage stem cell differentiation system such as the bone marrow hematopoietic system, the GRNs governing distinct cell fates may exhibit lineage-specific architecture. Thus, it would be desirable to facilitate lineage-specific GRN inference. The assignment of a particular cell to a lineage can be based on inferred cell fate probabilities, reflecting the likelihood of differentiating into any of the mature or terminal cell states present on the manifold. In the past, a number of computational methods for cell fate probability prediction has become available [1]. Modeling cell differentiation as probabilistic events allows us to regard this probability as a proximal measure of travel time from root cell states to terminal cell states, which we can connect with GRN construction through Granger regression modeling (Methods). This strategy enables the inference of cell fate- or lineage-specific GRNs.

To provide a proof-of-concept for integrating cell fate probabilities for lineage-specific GRN inference, we ran NetID on a hematopoietic cell state manifold comprising HSCs and lineage-biased progenitor cells [21]. We applied Palantir [17], which utilizes pseudotime information, to identify two dominating terminal cell states corresponding to the erythrocyte and the neutrophil lineage, and calculated the probability of each cell to transition to either state. Using a Granger causal model, we learned cell fate/lineage-specific networks governing erythroid and neutrophil differentiation.

Selection of the root cell is pivotal for pseudotime inference, since different root cells may lead to variability cell fate probability estimation. To resolve this issue and to increase stability of cell fate inference, Palantir or CellRank[27] allows to specify terminal states, which increases stability of cell fate inference. To test the robustness of lineage-specific GRN inference to root cell choice, we used different markers for root cell definition, or randomly sampled root cells from the multipotent progenitor population. After specifying the terminal states (neutrophils and erythroid cells), the inferred lineage-specific GRNs were very robust to root cell choice (Fig. S10).

In the Granger causal model, the number of coefficients that need to be estimated for each gene is *P* × *L. P* denotes the number of regulators and *L* denotes the maximum lagged time steps. To avoid overfitting, we applied L2 regularization to the Granger coefficients for each gene. We demonstrate this strategy by focusing on *KLF1* and the erythroid lineage as example target gene and lineage, respectively. After sorting the cells by cell fate probabilities, we splited data into training data (80%) and test data (20%). We trained the Granger causal model on training data and evaluated the prediction mean-squared error (MSE) and spearman correlation on test data. Without regularization (*λ* = 0), the MSE becomes larger and the spearman correlation decreases with increasing *P* and *L*. As the regularization strength *λ* is increased, the MSE is reduced and spearman correlation improves (Fig. S11A-B). Based on our trials we set *L* = 30 and *λ* = 150 as the default.

We next benchmarked NetID with alternative methods for lineage specific-GRN inference. We first conducted benchmarking on a simulated dataset with known ground truth lineage-specific GRNs (Methods). Using scVelo [19], a method that estimates the spliced RNA product velocity, on simulated data, we predicted the terminal states and the corresponding cell fate probabilities for the two lineages (CT1 and CT2, Fig. 5A-B).

**Figure 5.**
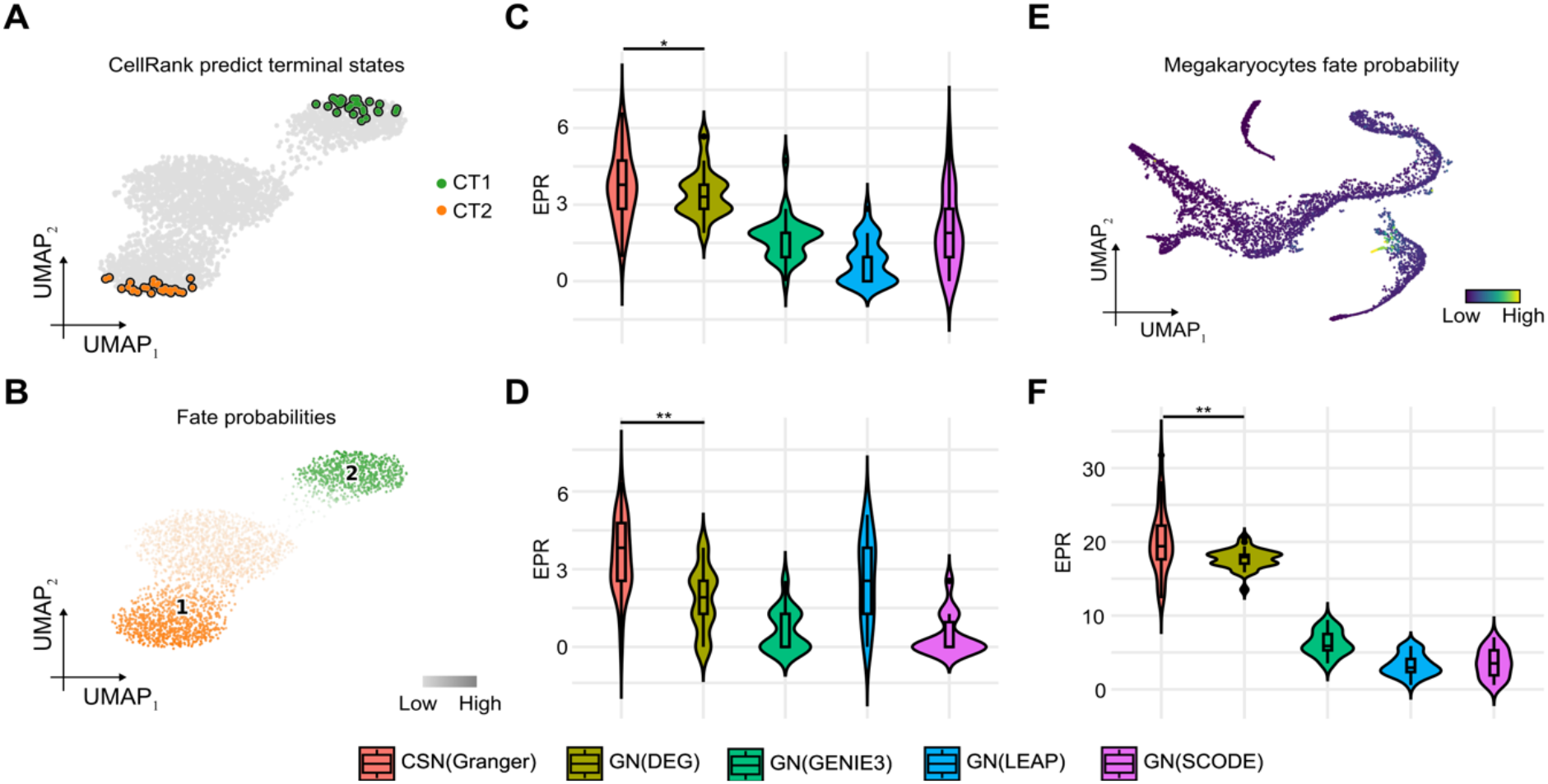
Incorporating cell fate probability improves GRN inference through NetID. A) UMAP plots of a dyngen simulated scRNA-seq dataset with two terminal states (CT1 and CT2). Each dot represents a cell, and the terminal states are colored. Terminal states are predicted using RNA velocity inferred by scVelo and cell fate prediction by CellRank. B) UMAP plots of a dyngen simulated scRNA-seq dataset with two terminal states (CT1 and CT2). Each dot represents a cell, and the terminal states are colored. The color scale indicates cell fate probability. C) Violinplot of EPR of the five methods. The simulated CT1 lineage-specific GRN was used as ground truth. Each method was run 30 times to derive the standard deviation. D) Violinplot of EPR of the five methods. The simulated CT2 lineage-specific GRN was used as ground truth. Each method was run 30 times to derive the standard deviation. D) UMAP plot of human hematopoietic differentiation dataset [17]. The 30 cells most confidently assigned to the terminal microstates are highlighted. E) Violinplot of EPR of the five methods. The GRN derived from a megakaryocyte-specific ChIP-seq dataset [30] was used as ground truth. Each method was run 30 times to derive the standard deviation. In (C, D and F), the box in the violinplot represents the interquartile range (IQR). The whiskers extend to the smallest and largest values within 1.5 times the IQR. The black line within the box indicates the median.. *P<0.05, **P<0.01, ***P<0.001, two-sided Wilcoxon rank sum test.

Using cell fate probability information for lineage-specific GRN inference significantly improved the performance in terms of AUPRC and EPR metrics (Fig. 5C-D, Fig. S12) compared with the global GRN estimated by GENIE3 and the GEINIE3-subnetwork derived from differential gene expression analysis (DEG) across lineages (Methods). Furthermore, NetID outperformed other network inference methods that directly incorporate pseudotime rather then cell fate probabilities, including SCODE [28] and LEAP [29] (Fig. 5C-D and Fig. S12).

For benchmarking on real data, we inferred the megacaryocytes-specific GRN for a human bone marrow dataset [17]. We first applied Palantir to predict megakaryocyte cell fate probabilities (Fig. 5E). We then inferred a megakaryocyte-specific GRN and compared to the GRN derived from a megakaryocyte-specific ChIP-seq dataset [30] as ground truth. Our results show that NetID with the Granger causal model outperforms all other methods. Although SCODE had a higher AUROC, its prediction precision was the lowest among the five methods (Fig. 5F and Fig. S12).

### NetID identifies key lineage-specific transcription factors and network modules

The benchmarking of NetID with different data preprocessing methods has demonstrated that using NetID-inferred metacell profiles preserves biological signals while also making GRN inference much more scalable. These characteristics suggest that NetID could enable GRN inference on large-scale scRNA-seq datasets. As a proof-of-principle, we applied NetID to a large mouse Kit+ hematopoietic progenitor scRNA-seq dataset with more than 40,000 cells comprising multiple lineages [31] (Fig. 6A). After running NetID, we found that metacell-derived gene expression profiles of transcription factors (TF) cluster into distinct modules, which were not detectable from raw gene expression values due to sparsity (Fig. S13A). Zooming in on specific pairs of TFs revealed that the covariation of metcell-based expression facilitates the recovery TF regulation from sparse data. For instance, we measured positive correlation between *Gata1* and *Klf1* (*R* = 0.875) or *Trp53* and *Pa2g4* (*R* = 0.795) (Fig. S13B), consistent with experimentally validated regulatory relationships [32, 33].

**Figure 6.**
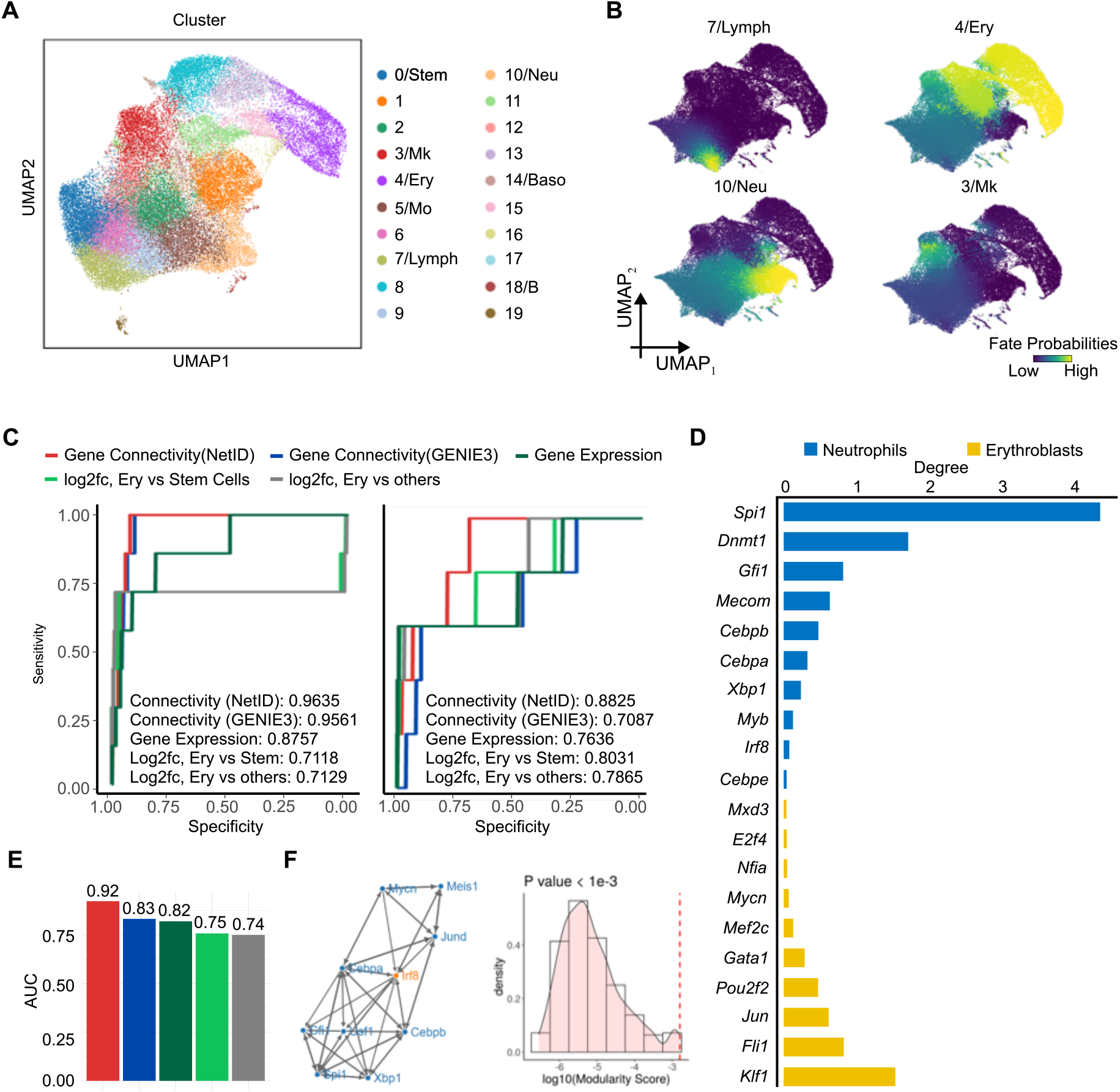
NetID infers lineage-specific GRNs in large-scale mouse hematopoiesis scRNA-seq data. A) UMAP plot of 40,000 mouse hematopoietic progenitor scRNA-seq data [31] colored by cell types. B) UMAP plot highlighting cell fate probabilities predicted by diffusion pseudotime analysis with Palantir. Cells are colored by the probability of giving rise to each of the four terminal states. C) Receiver operating characteristic (ROC) curve plots of five TF ranking methods: Gene Expression, log2-fold change (Log2fc) Ery vs Stem Cells, Log2fc Ery vs Others, and two network-based methods using the global network estimated by GENIE3 or NetID. Gene Expression: Ranking TFs according to the TF expression in both lineage terminal cell states. Log2fc, Ery vs Stem Cells: Ranking TFs according to the absolute log2-fold change of erythroid to stem cells. Log2fc, Ery vs others: Ranking TFs according to the absolute log2-foldchange of erythroid to other cells. D) Top 10 transcription factors (TFs) with the highest connectivity in lineage-specific gene regulatory networks inferred by NetID. F) Barplot of the average AUROC of the five TF ranking methods in (E). G) Visualization of the module centered on *Irf8* TF (left panel) and the histogram of the modularity score after 1000 permutations of the GRN structure. The red line indicates the modularity score of the original *Irf8* module (right panel). Stem: hematopoietic stem cells; Mk: megakaryocyte lineage; Ery: erythrocyte lineage; Neu: neutrophil lineage; Mo: monocyte lineage; Lymph: lymphocyte lineage; Baso: basophil lineage; B: B cell lineage;

As RNA velocity inference is known to be problematic for hematopoietic cell differentiation datasets, predicting inverted trajectories, we used Palantir to infer cell fate probabilities and identified four major terminal states in this dataset, corresponding to erythrocytes, megakaryocytes, neutrophils, and lymphocytes (Fig. 6B). To support the identification of key lineage-specific TFs by NetID, we calculated the regulator connectivity within each cell fate-specific network and ordered the TFs accordingly. The regulator connectivity reflects the importance of each gene in regulating other genes in the network. We found that the top-ranked TFs in the two main lineages, erythroid (Ery) and neutrophil (Neu), are distinct, and comprise many well-known regulators of the respective lineage (Fig. 6C). For instance, in the Ery branch, the top-ranked TF genes *Fli1* and *Klf1* are known as the master regulators of the megakaryocyte (Meg) and the Ery lineage, respectively [34]. The self-activation of FLI1 biases human stem or progenitor cells towards the Meg lineage, while KLF1 regulates differentiation towards the Ery lineage [34, 35]. The balance between Ery- and Meg-lineage cells is regulated by mutual antagonism of FLI1 and KLF1 [35]. For the Neu branch, the top ranking gene *Spi1* encodes the well-studied TF PU.1, a critical regulator of lymphoid-myeloid progenitor differentiation [36]. This TF and other key lineage-determining factors among the top-ranking TFs, such as *Cebpa*, are critical regulators of neutrophil differentiation [37, 38].

To systematically asses the performance of NetID in identifying key lineage factors compared to alternative methods, we used previously curated regulators of erythroid and neutrophil fate from the literature as ground truth [38-49] (Supplementary Table S2). We then computed the ranking of TFs based on regulator connectivity learned from NetID and other methods [35] and scored these rankings based on the ground truths (Fig. 6D). Several known TFs with critical functions during erythroid or neutrophil differentiation were among the top ranking genes. For the erythroid network, the top ranking gene *Klf1* is crucial for the expression of β-globin and other erythroid-specific genes [42]. Moreover, the eryrhoid lineage master regulator *Gata1*, which plays a pivotal role by activating erythroid-specific genes and repressing non-erythroid lineage genes [39] was recovered on rank 5.

For the neutrophil lineage, the top ranking gene was to *Spi1*, which encodes the master regulator PU.1 of myeloid lineage differentiation [38]. The high-ranking genes *Cebpa* encodes a known master regulator of granulocyte differentiation [50] and *Irf8* is critical in the early stages of neutrophil differentiation, where it acts alongside *Spi1* to drive the expression of myeloid-specific genes [46]. According to this benchmarking NetID outperforms all other methods, with an AUROC of 0.96 for the erythroid and 0.88 for the neutrophil lineage. Using one global network learned from GENIE3 only ranks Erythroid-associated TFs highly but shows reduced performance for the neutrophil lineage (AUROC of 0.73), suggesting that NetID’s inferred lineage-specific GRN captures lineage-specific features. Notably, we also found that using network-based ranking performs better than expression-based ranking, which supports the benefit of using gene regulatory network analysis for identifying lineage-specific regulators (Fig. 6E).

Zooming in on particular network module may reveal the regulatory underpinning of cell differentiation. By applying a permutation test (Methods) we could identify TF modules regulating differentiation. For instance, we identified a significant module centered on *Irf8* controlling neutrophil differentiation (*P* < 0.001, Fig. 6F). Further examination of this module revealed that *Irf8* interacts with *Cebpa* and *Cebpb*, forming a negative feedback loop with *Cebpb* [51] (Fig. 6F, Fig. S13). This feedback loop has been experimentally validated to control the chromatin state in dendritic cells. Additionally, we found that *Irf8* negatively regulates *Cebpa* (Fig. 6F, Fig. S14), in line with experimental evidence that Irf8 blocks the activity of *Cebpa* to prevent myeloid progenitor differentiation towards neutrophils [46]. We identified further examples of regulatory links predicted by NetID that have been validated experimentally in the past (Fig. S13).

## Discussion

GRN inference is a core objective of scRNA-seq data analysis for generating hypotheses on gene regulatory mechanisms underlying cell state transitions. However, GRN reconstruction for large scRNA-seq datasets is hampered by the requirement of computing gene-gene covariances across tens to hundreds of thousands of cells and scales with the square of the number of genes.

Moreover, gene expression quantification in individual cells is affected by substantial technical noise. Sampling of metacells or alternative imputation strategies represent potential solutions to this problem, but available methods suffer from the emergence of spurios gene-gene correlations not supported by the actual data [10]. To avoid such problems, it is critical to ensure cell state homogeneity within metacells and to make sure that each particular cell only contributes to a single metacell. Another key requirement is sufficient coverage of all cell states by metacells.

NetID addresses these challenges by optimizing metacell coverage of distinct states across the cell state manifold by geometric sketching [16], and ensures cell state homogeneity of each metacell by KNN graph pruning with VarID2. Overlap between distinct metacells is eliminated by a link probability-based reassignment strategy. Together, these steps permit scalable and accurate GRN inference and are not confounded by spurious correlations induced by common imputation strategies.

We validated the performance contribution of each step of the NetID algorithm using a hematopoietic progenitor differentiation dataset. By combining geosketch sampling, KNN graph pruning, and reassignment of partner cells, maximal GRN inference performance could be achieved. The optimal sample size for metacell inference was determined by balancing the number of sampled seed cells and the sparsity of the inferred metacell profile.

NetID was benchmarked against other imputation-based methods on simulated and real scRNA-seq datasets, and exhibited superior accuracy at substantially reduced runtime. Thus, NetID permits accurate GRN inference on large-scale scRNA-seq datasets, and overcomes scalability limitations of available methods.

We demonstrate that incorporation of cell fate probabilities enables lineage-specific GRN inference for multilineage scRNA-seq datasets. Our modeling approach, which relies on Granger causal regression enables the inference of directed regulator-target relationships, and was able to recover known TF network motifs driving differentiation of hematopoietic progenitors towards the erythrocyte and neutrophil lineages.

We note that future extensions of NetID could draw from previous approaches to overcome the issue of sparsity of scRNA-seq data for identifying lineage-specific regulators. In particular, incorporation of orthogonal datasets that are less affected by sparsity, such as bulk RNA- and ATAC-seq as well as ChiP-seq data, to infer regulon activity could help to prioritize key transcription factors driving lineage-specific GRN modules [52, 53].

In conclusion, NetD overcomes several limitations of currently available GRN inference approaches and provides a tool for interrogating the gene regulatory circuitry governing cell fate decisions in multilineage systems.

## Methods

### Inference of a pruned KNN network with VarID2

NetID requires the inference of a pruned KNN network from a single-cell gene expression matrix (*M* ∈ *R*^*n*×*m*^) through VarID2 (using the R package RaceID v0.3.1) [15] In short, VarID2 normalizes count data by negative binomial regression of total transcript counts, followed by principal component analysis (PCA) for initial dimensionality reduction. Subsequently, a fast KNN search is performed based on the Euclidean metric in PCA space to build the initial KNN network. For each cell in this network, VarID2 estimates the parameters of local negative binomial background distributions predicting the expected unique molecular identifier (UMI) count distribution for each gene. For cell *j*, the probability 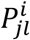 of the observed transcript count of gene *i* in each KNN *l* (*l* = *j*_1_, …, *j*_*k*_) is computed from this local distribution. The resulting link probabilities *P*_*jl*_ between cell *j* and each neighbor *l* are derived as the geometric mean of the count probabilities 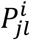 of the three genes *i* with lowest count probabilities. A larger probability *P*_*jl*_ indicates a stronger connection between cell *j* and its neighbor *l*. This calculation is performed for all links in the KNN network, to obtain a link probability matrix *P* ∈ *R*^*k*×*m*^ between each of the *m* cells and their *K* neighbors.

### Sampling seed cells from the transcriptomic manifold

As the next step of NetID, a representative subset of cells is sampled from the transcriptomic space with the objective that the sampled cells reflect the overall geometry of the dataset, and cover both rare and abundant cell stats. This strategy aims at preserving the transcriptional variation within the dataset and at the same time facilitates fast GRN computation. NetID implements two different sampling methods for dataset sketching: the first method is ‘geometric sketching’ (geosketch), which performs sketching on the principal component (PC) space. In the present analysis, we selected the top 50 PCs to reduce dimensions prior to running geosketch. The selected subset of sampled cells, termed “seed cells” is expected to capture the major transcriptome variation across the cell state manifold.

### Construction of metacells by aggregating pruned neighborhoods of seed cells

Conventional gene expression imputation methods rely on information sharing across neighborhoods in order to remove technical noise and achieve a more accurate estimate of gene expression for each cell. However, common imputation methods induce spurious correlations between the expression patterns of individual genes [12]. NetID relies on the inference of homogeneous, non-overlapping metacells. To achieve this, we first prune the neighborhood of each seed cell based on the link probability matrix *P* ∈ *R*^*k*×*m*^ inferred by VarID2 as described. The probabilities are compared to a probability threshold (*p*^*tr*^ = 0.01 by default), and all neighbors with *P*_*jl*_ < *p*^*tr*^ were pruned. Subsequently, remaining neighbors, termed partner cells, shared by different seed cells are assigned to the seed cell with the largest link probability. If the link probabilities with more than one seed cell are equal, the respective partner cell is assigned to the seed cell with the minimal number of neighbors to improve gene expression estimation accuracy. Finally, after all neighbors are pruned and assigned, we aggregate gene expression across all neighbors of each seed cell, and filter out the seed cells with too few partner cells (< 5 partner cells). This procedures yields the metacell gene expression profiles as input for GRN construction.

### Using non-linear methods to build basal GRN skeleton

Using the metacell gene expression profiles, NetID can accommodate any GRN construction method. According to a previous benchmarking study, PIDC [4], GENIE3 [2] and GRNBoost2 [54] were the top performing methods on real datasets. PIDC characterizes statistical dependencies between pairs of genes based on mutual information (MI), and the reliable estimation of pairwise joint probability distributions generally requires large sample size, i.e., a large number of cells. GENIE3 estimates gene-gene dependencies based on importance scores obtained be a random forests regression [55]. GRNBoost2 is based on the GENIE3 architecture, but utilizes gradient boosting. For consistency and comparability, we use GENIE3 (v1.20.0) to conduct GRN construction throughout the manuscript.

The output of these GRN construction methods is a matrix ***W*** of interaction coefficients between regulators and their predicted target genes. We further binarized the network by applying a threshold (0.001) on the weights and keeping the top *n* (default: 50) targets with the highest weight for each TF. To further improve the reliability of the predicted interactions, NetID allows intersecting the network skeleton with another network derived from bulk/single-cell ATAC-seq data obtained by combining predicted peak co-accessibility relationships and TF binding motif information. This prior skeleton matrix could be derived through scanning the open chromatin region within the promoter (±2*kb*) of each target. The correlation between the aggregated accessibility of this region and the target gene expression value could be used to filter out the regions that exhibit low correlation. A motif scanning method could then be used on these regions to identify the possible TFs that bind to it, to build a prior gene regulatory network skeleton from the epigenome dataset.

These GRN construction methods provide a non-linear view of global network structure in the single cell datasets, without considering lineage-specific architecture. Hence, we initially build a global GRN skeleton network, and then utilize inferred cell fate/lineage information to identify lineage-specific networks.

### Inferring lineage-specific GRNs from the global skeleton network through leveraging cell fate probabilities

Differentiation can be modelled as a probabilistic process. Existing approaches such as Palantir (v1.2) [17] model cell differentiation as a Markov process on KNN graphs inferred from transcriptome similarity. Alternative strategies rely on RNA velocity estimation [18] to infer vector fields representing differentiation trajectories on cell state manifolds. CellRank (v1.5.2) [27] combines similarity based Markov models with RNA velocity to enhance cell fate probability estimation. In general, these methods predict each cell’s probability to differentiate towards any of the mature fates, or lineages, in the system. We leveraged cell fate probability predictions obtained by these approaches to build cell fate specific GRNs from the skeleton network.

NetID integrates CellRank or Palantir for cell fate probability prediction. These methods output a cell fate matrix *F* ∈ *R*^*k*×*m*^, where each row represents a cell and each column represent a cell fate (or terminal state) and the entries correspond to cell fate probabilities. First, cells are assigned to fates/lineages in the manifold through clustering based on cell fate probabilities. Specifically, we applied a Gaussian Mixture Model with an optimal cluster number selected by Bayesian-Information-Criterion (BIC) to cluster cells using the cell fate probability matrix. A cluster is then assigned to a given lineage if the mean cell fate probability of the cluster towards this lineage is *k*-fold (k=2 by default) higher than for any other lineage. Clusters with comparable cell fate probabilities towards all lineages are regarded as uncertain states. For the clusters with comparable cell fate probabilities towards multiple lineage, we assign the cluster to the lineages they are biased to. Each cell lineage is defined as the union of cells in clusters assigned to that lineage and the uncertain states.

For each lineage *k*, we ordered the metacells (the metacells we use to construct the global GRN In the last step) according to their average cell fate probability, yielding time series of *N* genes with timestamps *t* = 1,2, …, *T* where *T* corresponds to the number of metacells in the lineage. The expression of gene *i* at timepoint *t* is denoted as *x*_*i*_(*t*). Using the global skeleton network to determine the target and regulator genes, we can build a Granger causal regression model for each target gene *i* through minimizing the loss function

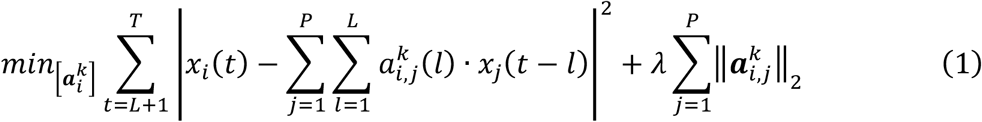

where 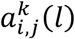 corresponds to the *l*-th lagged Granger coefficient from the time series of regulator genes *x*_*j*_ to the time series of the target gene *x*_*i*_ in lineage *k. P* denotes the number of regulators of target gene *i. L* denotes the maximum lagged time steps. The estimated *P* × *L* coefficient matrix 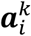 represents the Granger coefficients governing the time series of gene expression readout. *λ* denotes the regularization parameter. In this studies, we use *L* = 30 and *λ* = 150 to conduct all Granger regression analyses.

Furthermore, we quantify the edge weight of regulator gene *j* to target gene *i* for lineage *k* as follow:

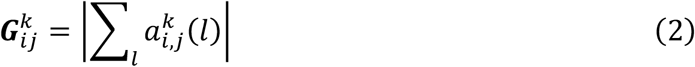

The output weight matrix ***G***^***k***^ represents the Granger score based GRN learned for lineage *k*. We rank-transformed the interaction weights between a regulator and its targets. More precisely, we transformed each row of ***G***^***k***^, by replacing each interaction weight by 1/*i*^2^ where *i* denotes the rank of the interaction weight between a particular regulator and its targets ordered by decreasing interaction weights. Entries for non-interacting genes are replaced by zeroes. The final interaction weights represents the learned lineage-specific GRN 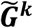.

**Identification of the optimal number of sampled seed cells**

Analyzing the tradeoff between the number of sampled seed cells and the resulting effective size of metacells (the number of partner cells) allows us to determine an optimal sampling size. Suppose *S* is the sample size, *n*_*s*_ is the average effective size for metacells (average number of neighborhoods across all seed cells) given a specific sampling size *S*. We vary *S* from 50 to 1000 cells (or up to all cells). The resulting score for sample size *S* is defined as follow:

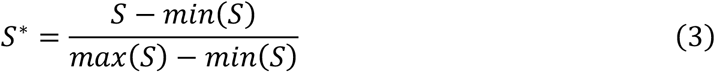

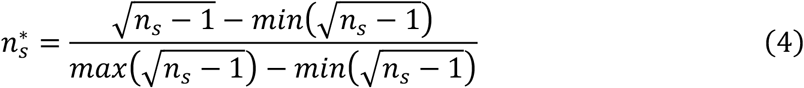

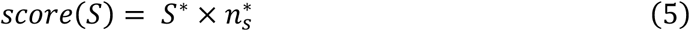

The optimal sample size is defined as:

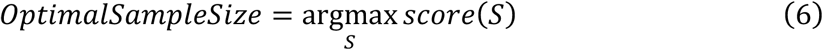

### Metrics for GRN benchmark

#### Explained Expression Variance

A good cell sampling method maximizes the recovered variance of the original data at a given sample size. We define the *Explained Expression Variance* metric (EEV) to benchmark different sampling methods according to this objective: First, we apply PCA to decompose the original expression matrix into the top *K* (*K*= 10 by default) principal components (PCs). For each PC, we evaluate a regression model with the sampled cells as predictors, and calculate the goodness-of-fit (R^2^). The EEV value is define as follow:

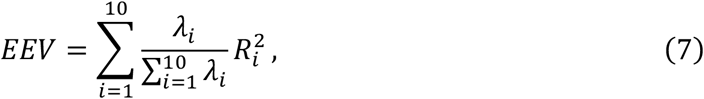

Where *λ*_*i*_ is the eigenvalue of i-th PC.

#### Early Precision Rate (EPR)

Early precision is defined as the fraction of true positives in the top *k* edges (*k*=the number of edges in the ground truth network by default) where the top *k* edges are regarded as the positive prediction. Then the early precision rate (EPR) represents the ratio of the early precision value and the early precision for a random predictor for this network. A random predictor’s precision is the edge density of the ground-truth network. The EPR measures how well an algorithm is able identify true positive interactions early on in the ranking. A detailed explanation of the EPR is shown as conceptual Fig. S1.

#### AUPRC, AUROC

We computed areas under the precision-recall and receiver operating characteristic curves using the edges in the respective ground truth network ranked edges from each method as the predictions.

Both AUPRC and EPR are metrics to evaluate precision. However, different to AUPRC, which considers all predictions in the ranking, the EPR focuses on the early predictions only. In experimental single-cell RNA-seq data, the number of cells and the expression variability can vary widely across genes, which can affect the performance of GRN inference algorithms. The EPR metric may be more robust to such variability, as it focuses on the top predictions that are most likely to be biologically relevant.

Therefore, in simulated datasets with known ground truth regulatory network, we computed EPR, AUPRC, and AUROC for benchmarking. For real datasets, we used ChIP-seq or STRING networks as the proxy for ground truth and only computed EPR for benchmarking.

### Simulated datasets for benchmarking

We utilized dyngen [24] to conduct simulations of gene expression in single cells. Specifically, we generated two separate single cell gene expression manifolds featuring either a cycling or bifurcating topology (Fig. 3). These datasets were chosen specifically to be used as benchmarks for the evaluation of gene regulatory networks (GRNs). Each dataset simulates 50 TFs, 200 targets and 50 house keeping genes, in 4,000 cells. For all other parameters we used the default setting of dyngen.

In Fig. 5, we utilized dyngen to simulate bifurcating topology scRNA-seq data with cell-specific ground truth GRNs. To define lineage-specific GRNs, we aggregated all cell-specific GRNs for lineages CT1 or CT2, respectively. To obtain the aggregated GRN for a cell type, e.g. CT1, we sum up the cell-specific network of all cells belonging to this lineage:

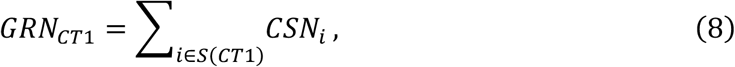

*S*(*CT*1) denotes the set of all cells belonging to the CT1 terminal states, and *CSN*_*i*_denotes the cell-specific network of cell *i*.

See https://github.com/WWXkenmo/NetID_package/blob/main/dyngen_simulation_netID.r for the code of the dyngen simulations.

### Lineage specific GRN inference through DEG analysis

Assuming that the GENIE3-inferred global network already contains all lineage-specific regulatory information, one effective strategy for extracting lineage-specific GRNs is through differential gene expression (DEG) analysis. First, we identify the highly up-regulated genes in each terminal state using the Wilcoxon rank-sum test. We keep only those genes with an adjusted p-value < 0.05 in the GENIE3 network, thereby deriving a lineage-specific GRN for each terminal state.

### GRN inference and ground truth GRN in real datasets

To benchmark GRN inference performance for real datasets, after processing of raw gene expression data according to each method, we applied the same GRN inference algorithm, GENIE3, to build a GRN for each method. We only constructed the GRNs for the TFs in top 3,000 highly variable genes. The TF list was curated from the BEELINE pipeline (https://github.com/Murali-group/Beeline) [3].

For each dataset, we selected two categories of biological networks from BEELINE as ground truth. The first category is a general network with no contextual specificity, which includes non-specific tissue ChIP-seq networks and STRING [56] functional networks. The non-specific ChIP-seq networks were extracted from three resources: DoRothEA [57], RegNetwork [58] and TTRUST [59]. In DoRothEA, we only considered two levels of evidence: A (curated/high confidence) and B (likely confidence).

The second category is the cell type-specific network. For the mESC dataset, we selected loss-of-function/gain-of-function (lof/gof) networks and mESC-specific ChIP-seq networks, while for the mHSC dataset, we selected mHSC-specific ChIP-seq networks. Those cell type specific ChIP-seq networkds were extracted from three sources: ENCODE[60], ChIP-Atlas [61] and ESCAPE[62].

The scRNA-seq datasets and ground truth GRN resources used in this work are provided in Supplementary Table S1.

### Cell fate prediction with Palantir or CellRank

Palantir and CellRank are used for predicting cell fate probability. When applying Palantir to mouse hematopoietic data from Tusi et al. [21], we specified the cell with the highest *Runx2* expression as root cell. When applying Palantir on the bone marrow dataset from Setty et al. [17], we used precomputed Palantir pseudotime and the terminal states specified as “CLP”, “Mono 1”, “DCs”, “Ery_2” and “Mega”. When applying Palantir to mouse hematopoietic data from Dahlin et al. [31], we selected the cell with the highest *Procr* expression as the root cell and specified the terminal states as “7/Lymph”, “4/Ery”, “10/Neu” and “3/Mk”. When applying CellRank on the simulated dataset [63], we used the dynamical model to calculate velocity in scVelo.

### Regulatory Module Identification and statistics significant test

First, we ranked the transcription factors (TFs) by their gene connectivity, which is defined as the sum of the regulatory coefficients of each TF. We ordered the connectivity values to derive the top *k* most important TFs. Using these TFs as seeds, we ran the spinglass clustering algorithm [64] on the gene regulatory network (GRN) to identify the module closest to these TFs.

To evaluate the statistical significance of each module, we permuted the gene IDs of the GRN 1,000 times, resulting in 1,000 different permuted GRNs with the same topological structure as the original network. For each module, we calculated the modularity score, which is defined as the average regulatory coefficient within that module, for both the original network and each of the randomized networks. We then calculated the permutation p-value for each module. The permutation p-value is calculated by comparing the modularity score of the module in the original network with the distribution of modularity scores obtained from the randomized networks.

## Data availability

The following publicly available datasets were used in this study:

Mouse hematopoietic dataset from Tusi et al. [21] (GSE89754)

Human adult hematopoietic dataset from Buenrostro et al. [22] (GSE96772)

Human bone marrow dataset from Setty et al. (https://explore.data.humancellatlas.org/projects/091cf39b-01bc-42e5-9437-f419a66c8a45)

Mouse hematopoietic stem cell dataset from Pei et al. [25] (GSE152555)

Mouse embryonic stem cell dataset from Klein et al. [23] (GSE65525)

Mouse embryogenesis dataset from Qiu et al. [26] (http://tome.gs.washington.edu)

Mouse pancreatic single cell dataset from Bastidas-Ponce et al. [63] (GSE132188)

Mouse hematopoietic dataset from Dahlin et al. [31] (GSE107727)

## Code availability

The NetID open-source code is maintained and documented on GitHub (https://github.com/WWXkenmo/NetID_package) and is publicly available under the MIT license.

## Acknowledgements

We extend our heartfelt gratitude to Professor Ting Ni for his support and contributions to this study. This work was primarily conducted in Professor Ni’s laboratory, and his generous provision of computational resources played a pivotal role in the successful execution of our research.

## Author contributions

DG conceived the study. WW and DG designed the algorithm. WW implemented the algorithm and performed most of the computational analyses under supervision of DG. YW and RL performed the methods benchmark, WW and DG wrote the paper.

## Supplementary Information

Additional file 1: Supplementary Table S1-S2, Supplementary Figures S1-S14.

