## Supplementary Figures for "Scalable identification of lineage-specific gene regulatory networks from metacells with NetID"

### Supplementary Material

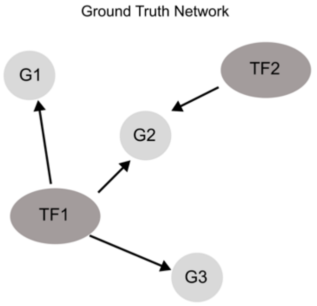


GENIE3 network prediction

| Source | Target | True Links | Rank Score |
|---|---|---|---|
| TF1 | G1 | 1 | 0.9 |
| TF2 | G1 | 0 | 0.45 |
| TF1 | G3 | 1 | 0.3 |
| TF2 | G2 | 1 | 0.1 |
| TF2 | G3 | 0 | 0.05 |
| TF1 | G2 | 1 | 0.01 |

Top *K* (first four rows)

$$EarlyPrecision = \frac{Hits@K}{K}$$

$$NetworkDensity = \frac{K}{MP}$$

$$EPR = \frac{EarlyPrecision}{NetworkDensity}$$

*M* : Number of targets
*P* : Number of regulators
*K* : Number of edges

**Figure S1. Conceptual figure for early precision rate (EPR).**
Early Precision Rate (EPR) is a metric proposed in BEELINE [1] to evaluate early precision compared to a random predictor. EPR is defined as the fraction of true positives among the top-k edges, where k is the number of edges in the ground-truth network. In our benchmark, we consider all predicted top-k edges as positive for all methods, making EPR conceptually similar to precision, which is the fraction of true positive instances among all positive instances. In this figure, we present a toy example with 2 transcription factors (TFs) and 3 targets. In this case, following the described method, EPR is calculated as (3/4) / (4/6) = 1.125.

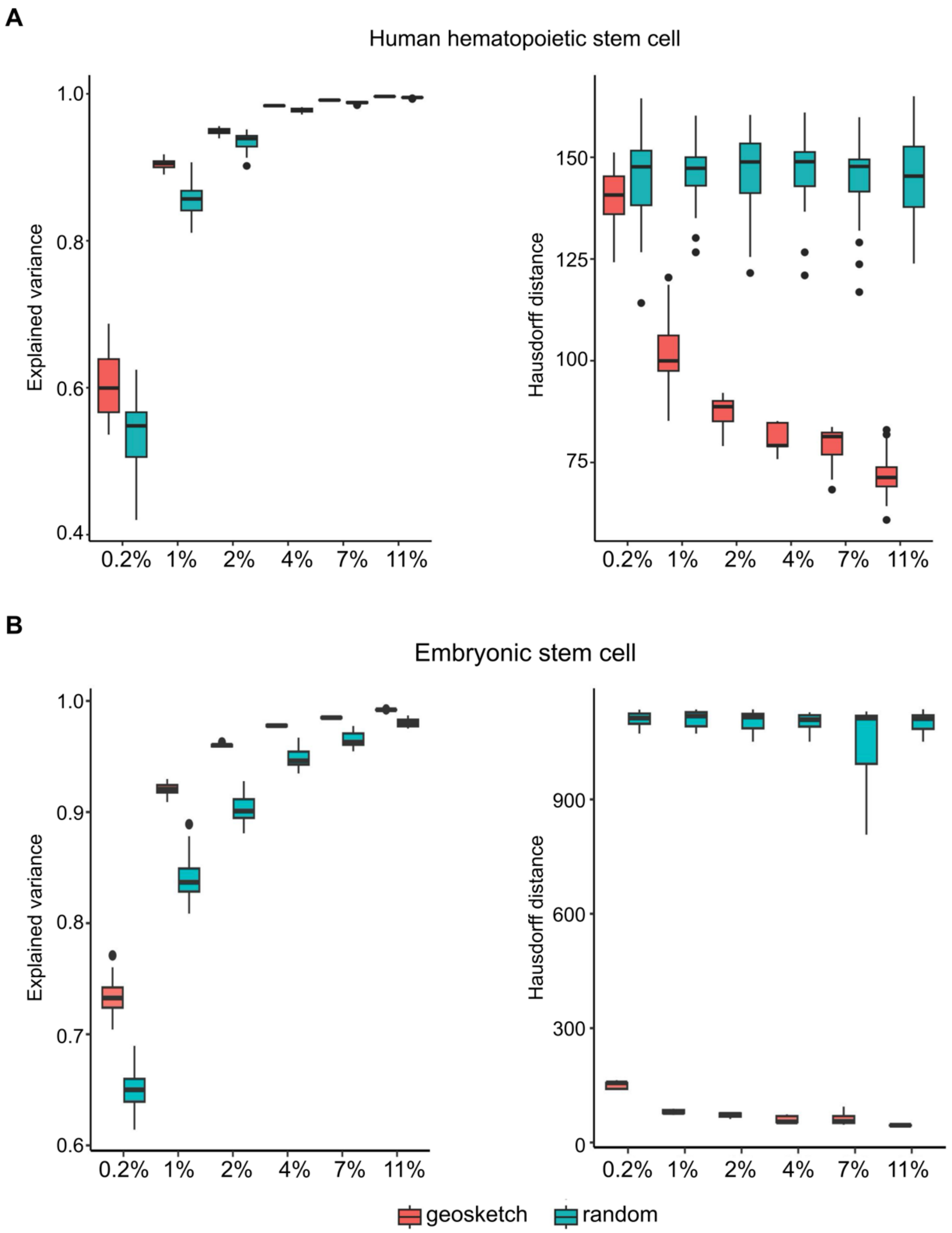


**Figure S2. Comparing geosketch and random sampling effects.**

Comparing different sample model effects to the human adult hematopoietic differentiation [2] and embryonic stem cell dataset [3].

A. Boxplot comparing the Hausdorff distance (left) and Explained Expression Variance (right) of randomly sampled cells (green) and geosketch sampled cells (red) with 30 repeats in human hematopoietic stem cell dataset. The x-axis denotes the number of sampled cells.

B. Boxplot comparing the Hausdorff distance (left) and Explained Expression Variance (right) of randomly sampled cells (green) and geosketch sampled cells (red) with 30 repeats in mouse embryonic stem cell dataset. The x-axis denotes the number of sampled cells.

In (A-B), the box in the boxplot represents the interquartile range (IQR).The whiskers extend to the smallest and largest values within 1.5 times the IQR. The black line within the box indicates the median.

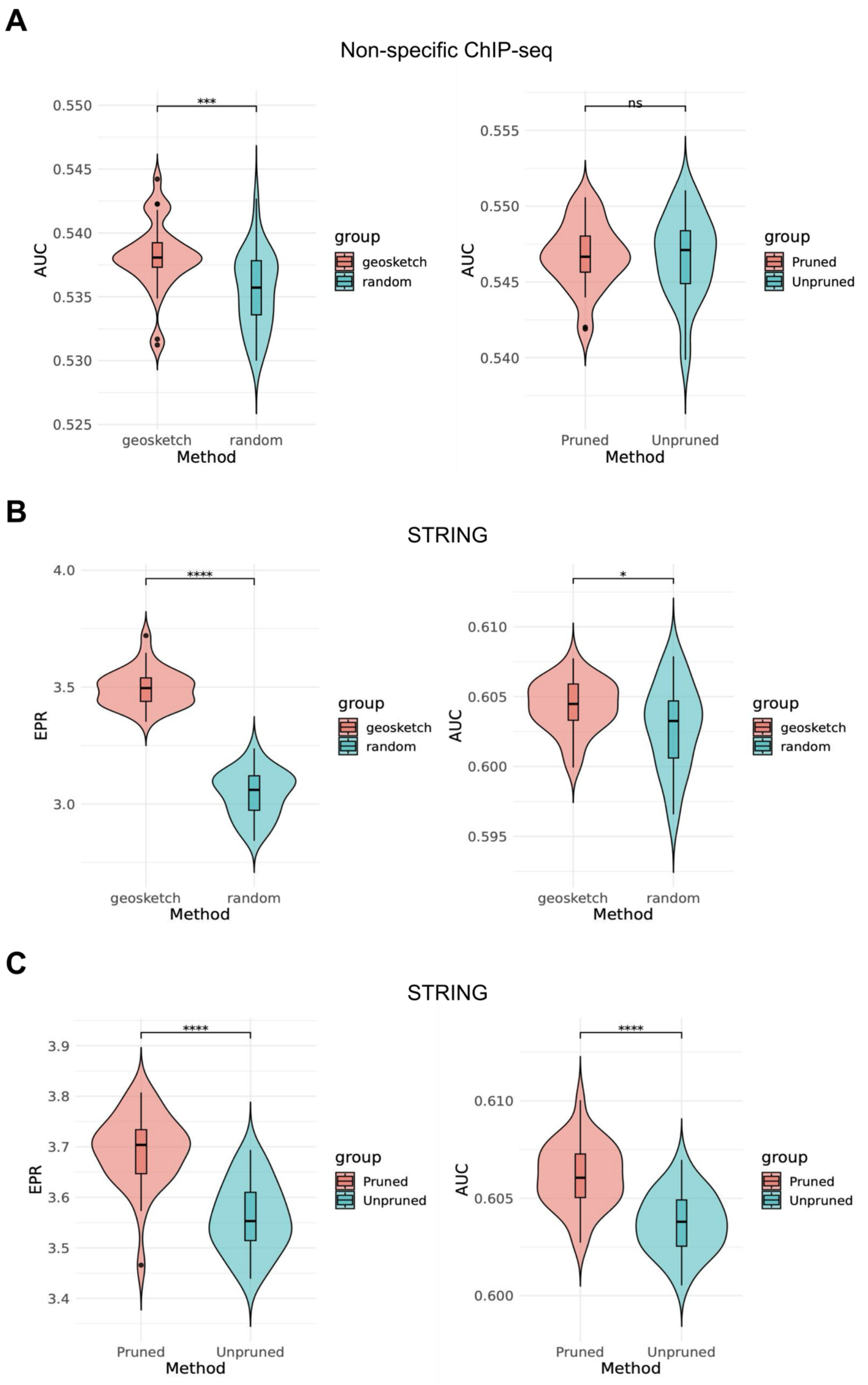


**Figure S3. Comparison of inferred GRN using different sampling strategies and**

**graph pruning in mouse hematopoietic differentiation.**

A. Violinplot showing the difference in Area Under the Receiver Operating Characteristic Curve (AUROC) when using GENIE3 inferred GRN on geosketch sampled cells or randomly sampled cells (left) and pruned KNN graph or unpruned KNN graph (right). We used non-specific ChIP-seq dataset as ground truth.

B. Violinplot showing the difference in EPR (left) and AUROC (right) when using GENIE3 inferred GRN on geosketch sampled cells or randomly sampled cells. We used STRING dataset as ground truth.

C. Violin-plot showing the difference in EPR (left) and AUROC (right) when using GENIE3 inferred GRN on pruned KNN graph or unpruned KNN graph. We used STRING dataset as ground truth.

In (A-C), the box in the violinplot represents the interquartile range (IQR).The whiskers extend to the smallest and largest values within 1.5 times the IQR. The black line within the box indicates the median.

* P<0.05, ** P<0.01, *** P<0.001, two-sided Wilcoxon rank sum test.

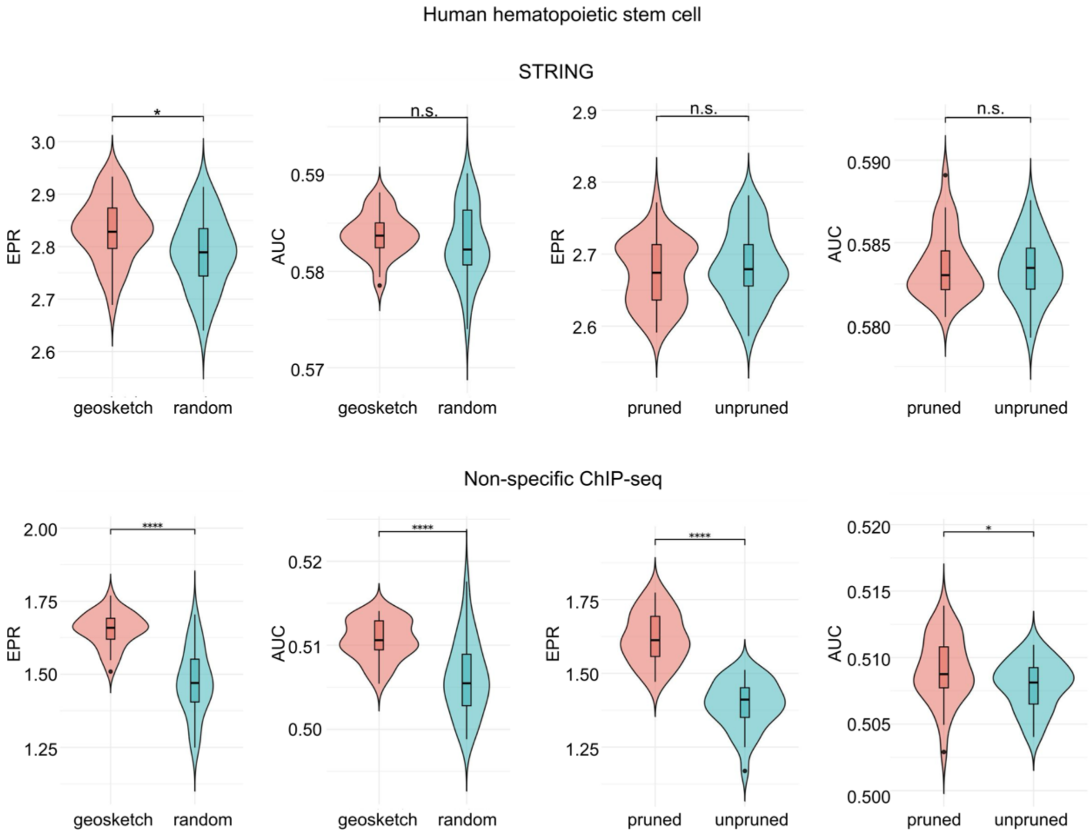


**Figure S4. Benchmark GRN step-by-step to comparing the effects of sampling and pruning in human adult hematopoietic differentiation [2].**

We benchmarked sampling (geosketch vs. random) and pruning (pruned vs. unpruned) effects on GRN construction step-by-step. Data are presented as violin plots for both non-specific ChIP-seq data (top) and STRING datasets (bottom) as the ground truth.

The box in the violinplot represents the interquartile range (IQR).The whiskers extend to the smallest and largest values within 1.5 times the IQR. The black line within the box indicates the median.

* P<0.05, ** P<0.01, *** P<0.001, two-sided Wilcoxon rank sum test.

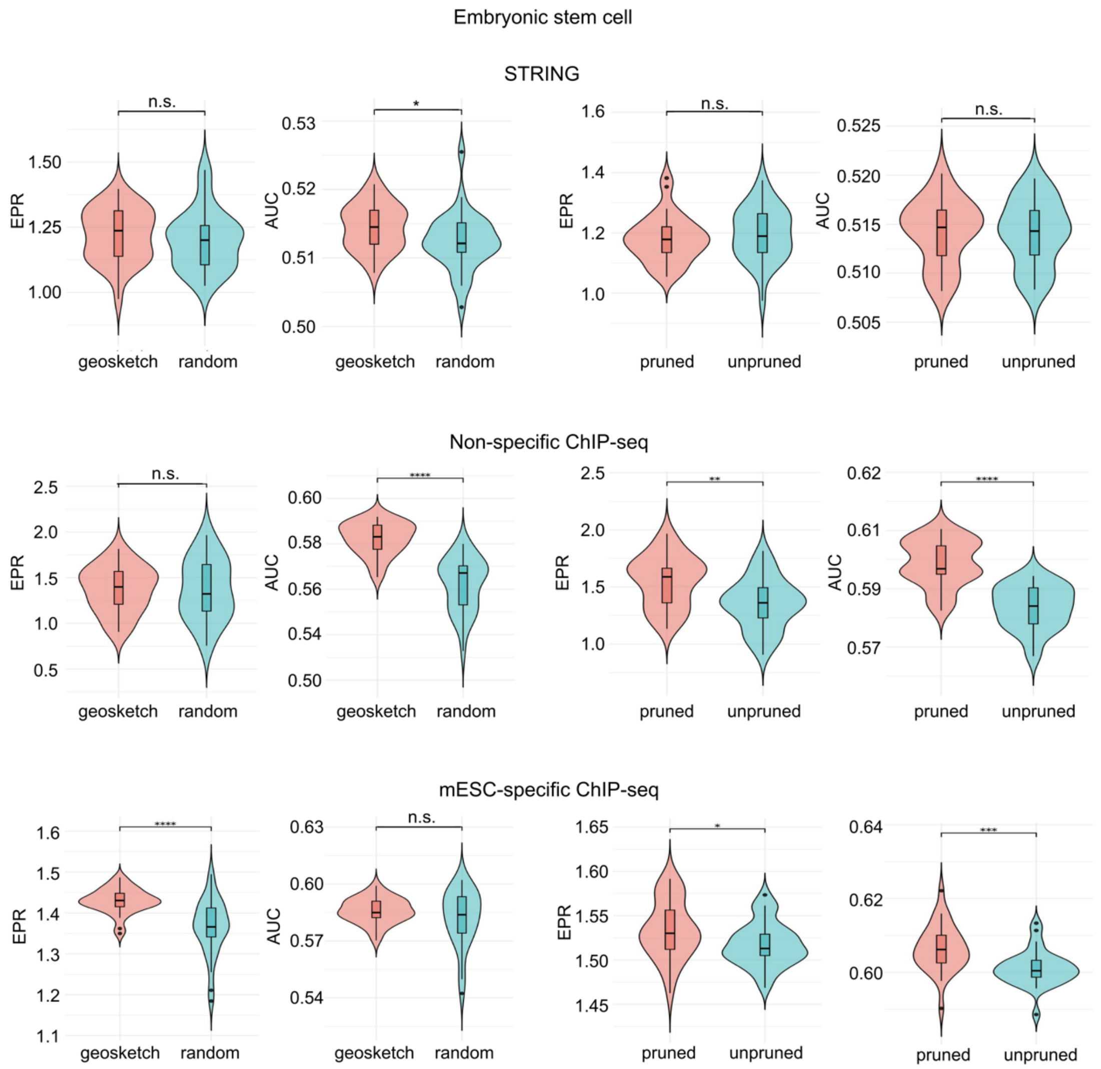


**Figure S5. Benchmark GRN step-by-step to comparing the effects of sampling and pruning in mouse embryonic stem cell (mESC) dataset [3].**

We benchmarked sampling (geosketch vs. random) and pruning (pruned vs. unpruned) effects on GRN construction step-by-step. Data are presented as violin plots for STRING datasets (top), non-specific ChIP-seq data (middle) and mESC-specific ChIP-seq data datasets (bottom) as the ground truth.

The box in the violinplot represents the interquartile range (IQR).The whiskers extend to the smallest and largest values within 1.5 times the IQR. The black line within the box indicates the median.

* P<0.05, ** P<0.01, *** P<0.001, two-sided Wilcoxon rank sum test.

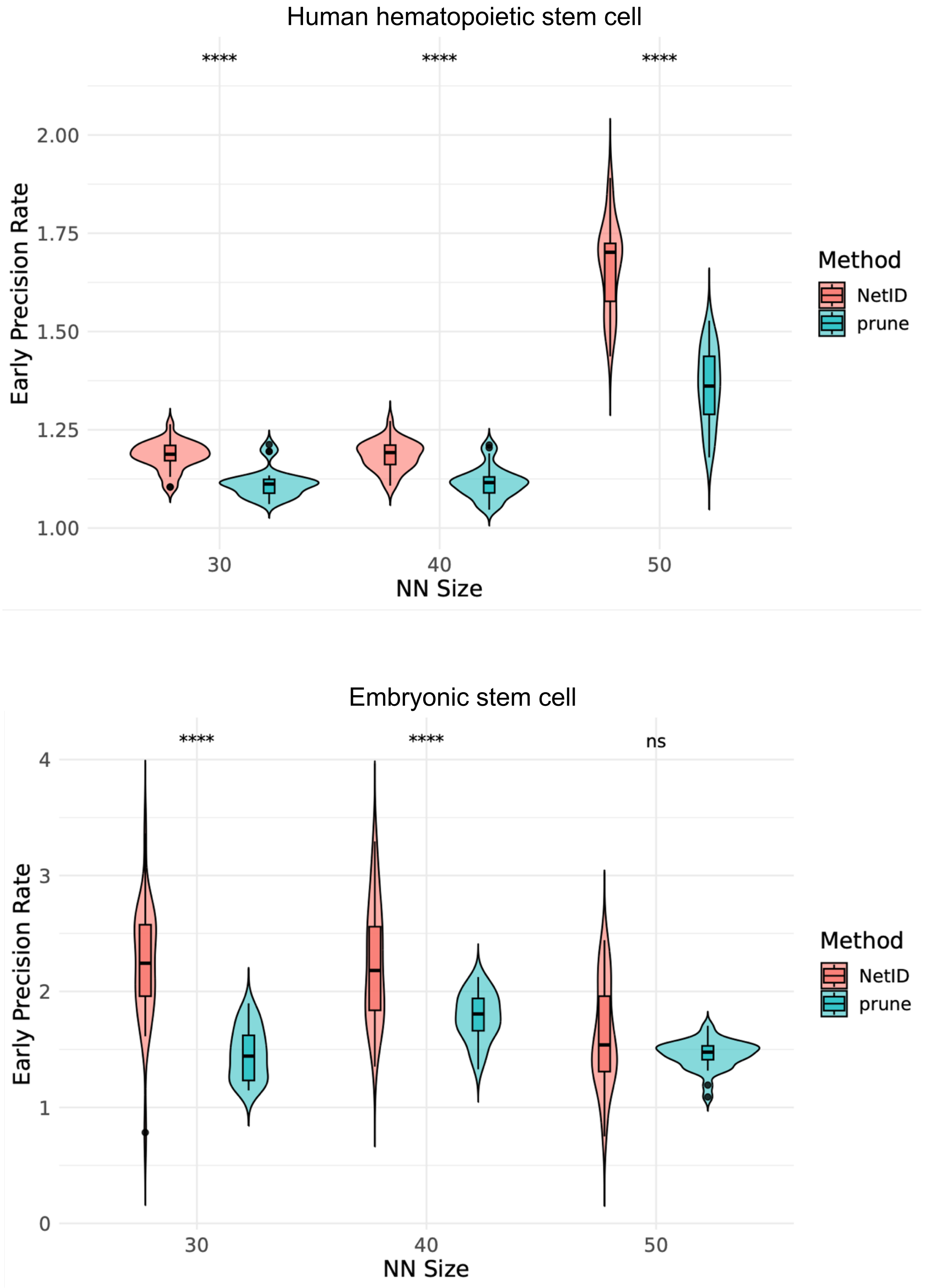


**Figure S6. Comparing the effect of neighbor reassignment in the human adult hematopoietic differentiation and mouse embryonic stem cell (mESC) dataset.** Violin-plots showing the comparison of early precision rate between two different strategies. "NetID" denotes combining geosketch, KNN graph pruning, and neighbor reassignments to build metacell profiles for GRN inference with 30 repeats. "Prune"

denotes KNN graph pruning, and neighbor reassignments to build metacell profiles for GRN inference with 30 repeats. The x-axis denotes the number of neighbors. We benchmarked the performance on two biological networks as ground truth: the non-specific ChIP-seq network (human adult hematopoietic) and mESC specific ChIP-seq network (mESC dataset).

The box in the violinplot represents the interquartile range (IQR).The whiskers extend to the smallest and largest values within 1.5 times the IQR. The black line within the box indicates the median.

* P<0.05, ** P<0.01, *** P<0.001, two-sided Wilcoxon rank sum test.

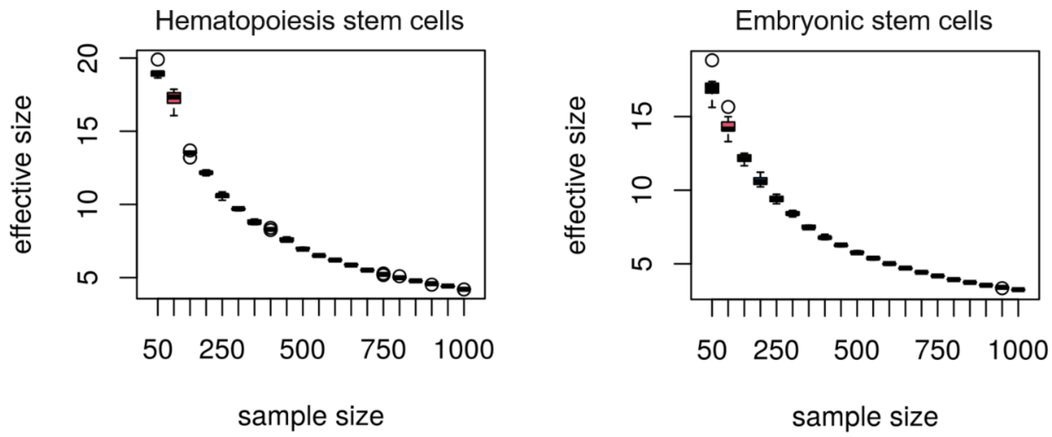


**Figure S7. Distribution of the number of partner cells per seed cell in different sample size**

Boxplot of the number of sampled seed cells' partner cells. The x-axis denotes the number of sampled seed cells. The y-axis denotes the number of partner cells per seed cells.

The box in the boxplot represents the interquartile range (IQR).The whiskers extend to the smallest and largest values within 1.5 times the IQR. The black line within the box indicates the median.

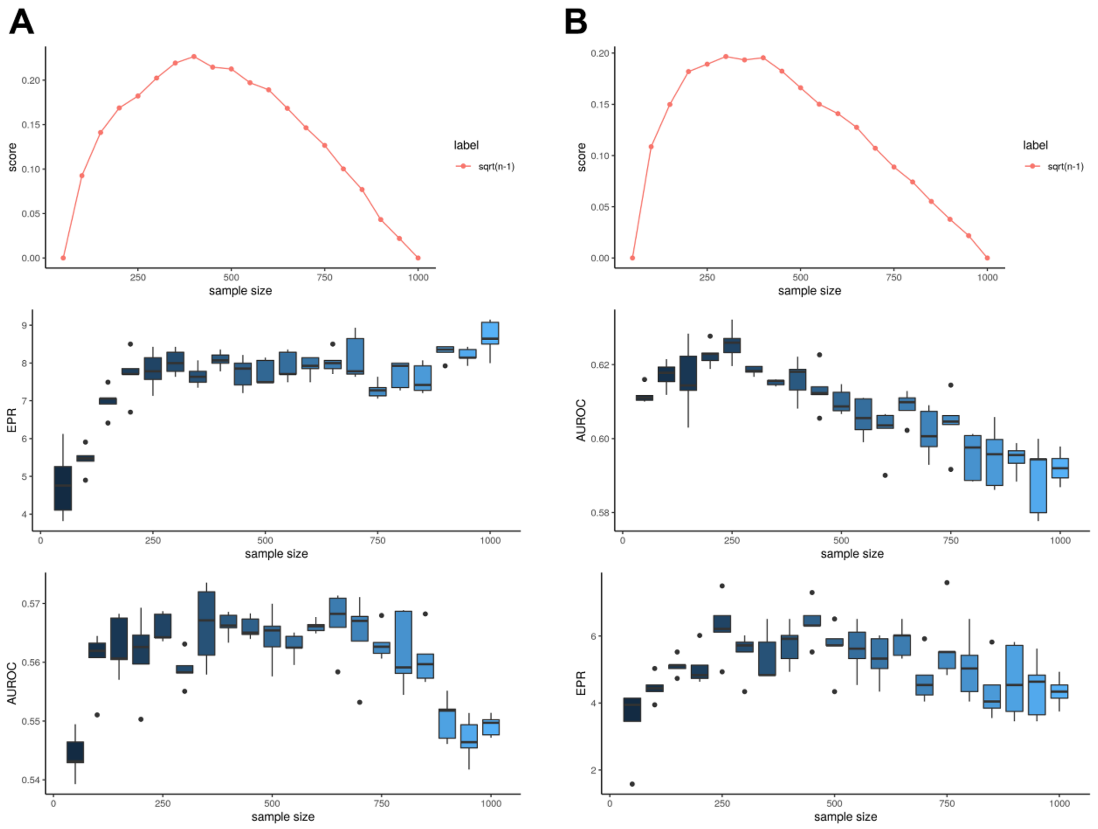


**Figure S8. Optimizing the number of sampled seed cells.**

A. Lineplot showing the sample size score as a function of sample size (top). The boxplots show the difference in Early Precision Rate (EPR) (middle) and Area Under the Receiver Operating Characteristic Curve (AUROC) (down) for GENIE3-inferred GRNs on different numbers of sampled seed cells. Data are shown for the mouse hematopoiesis stem cell gene expression dataset and the non-specific ChIP-seq network was used as the ground truth.

B. Lineplot showing the sample size score as a function of sample size (top). The boxplots show the difference in Early Precision Rate (EPR) (middle) and Area Under the Receiver Operating Characteristic Curve (AUROC) (down) for GENIE3-inferred GRNs on different numbers of sampled seed cells. Data are shown for the mouse embryonic stem cell gene expression dataset and the non-specific ChIP-seq network was used as the ground truth.

The box in the boxplot represents the interquartile range (IQR).The whiskers extend to the smallest and largest values within 1.5 times the IQR. The black line within the box indicates the median.

* $P<0.05$, ** $P<0.01$, *** $P<0.001$, two-sided Wilcoxon rank sum test.

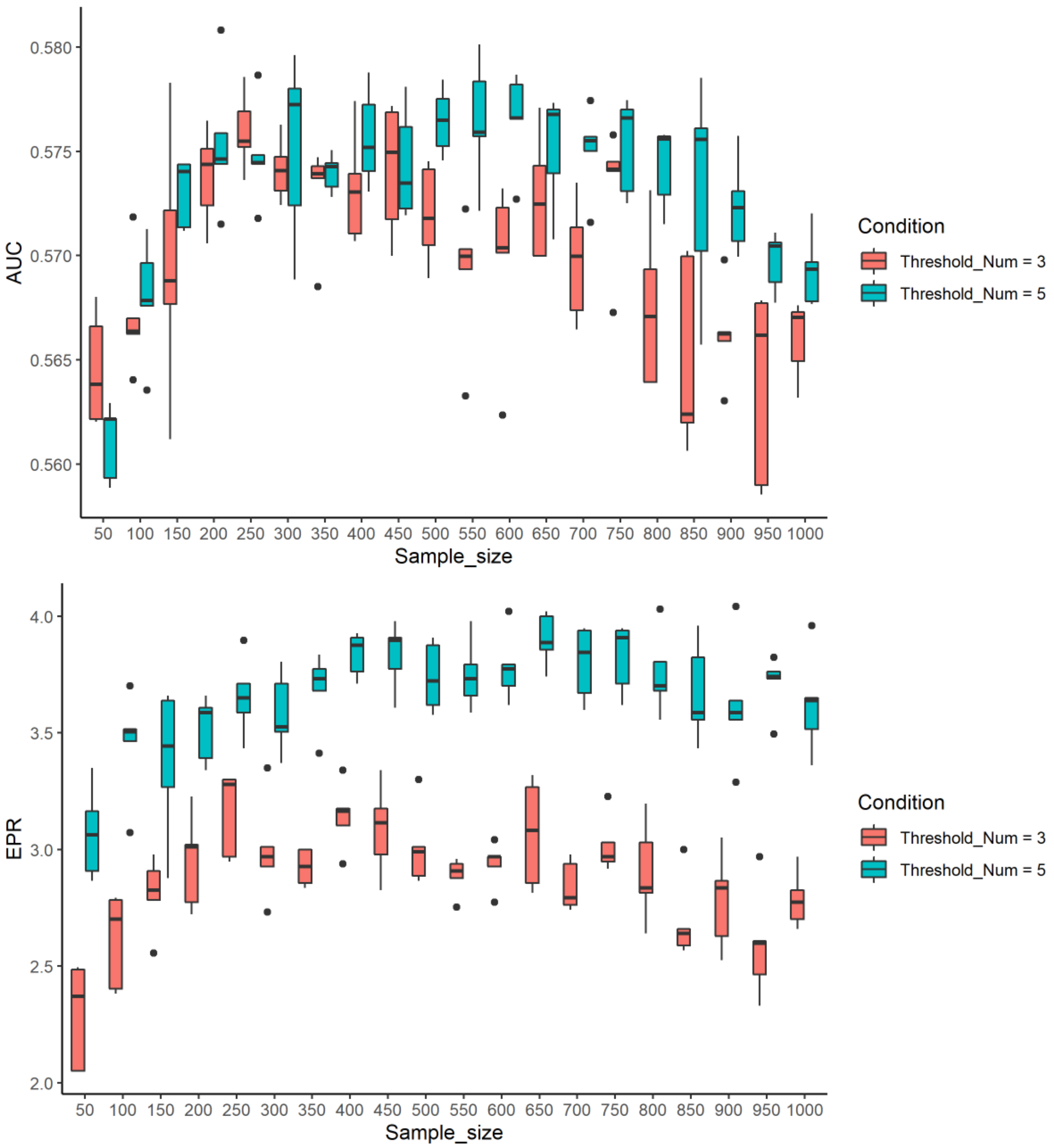


**Figure S9. Seed cells filtering effects on GRN inference**

Boxplots showing the difference in Area Under the Receiver Operating Characteristic Curve (AUROC) and Early Precision Rate (EPR) for different threshold numbers to filter seed cells. We used the mouse hematopoiesic stem cell gene expression dataset for GRN inference and the non-specific ChIP-seq network as the ground truth.

The box in the boxplot represents the interquartile range (IQR).The whiskers extend to the smallest and largest values within 1.5 times the IQR. The black line within the box indicates the median.

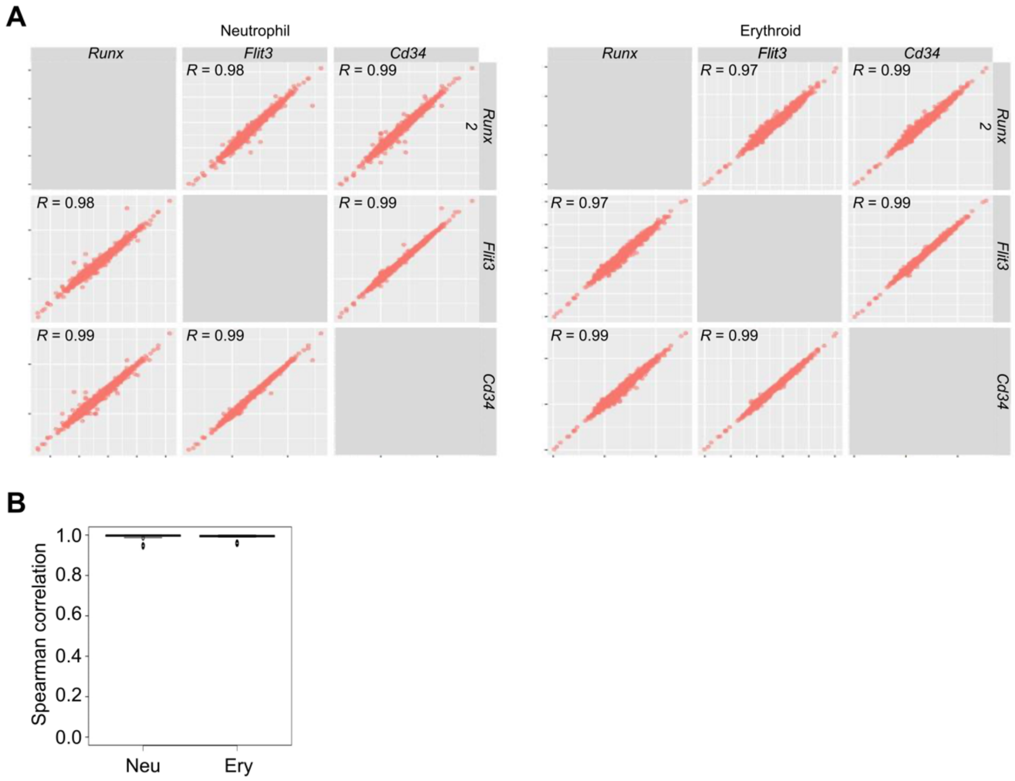


**Figure S10. Robustness of GRN inference to root cell selection.**

A. Pairwise scatter plots display the comparison of lineage-specific gene regulatory networks (GRNs) for the mouse hematopoietic differentiation dataset. We applied NetID for GRN prediction, with cell fate determined by Palantir. Hematopoietic stem cell and progenitor markers *Runx2*, *Flt3*, and *Cd34* were used to infer cell fate probabilities. The resulting lineage-specific GRNs were compared, specifying terminal states as neutrophils and erythroid cells.

B. Boxplots display the comparison of lineage-specific gene regulatory networks (GRNs) for the mouse hematopoietic differentiation dataset. We applied NetID for GRN prediction, with cell fate determined by Palantir. Randomly sampled cells from multipotent progenitor cells were used to infer cell fate probabilities. The resulting lineage-specific GRNs were compared, specifying terminal states as neutrophils and erythroid cells.

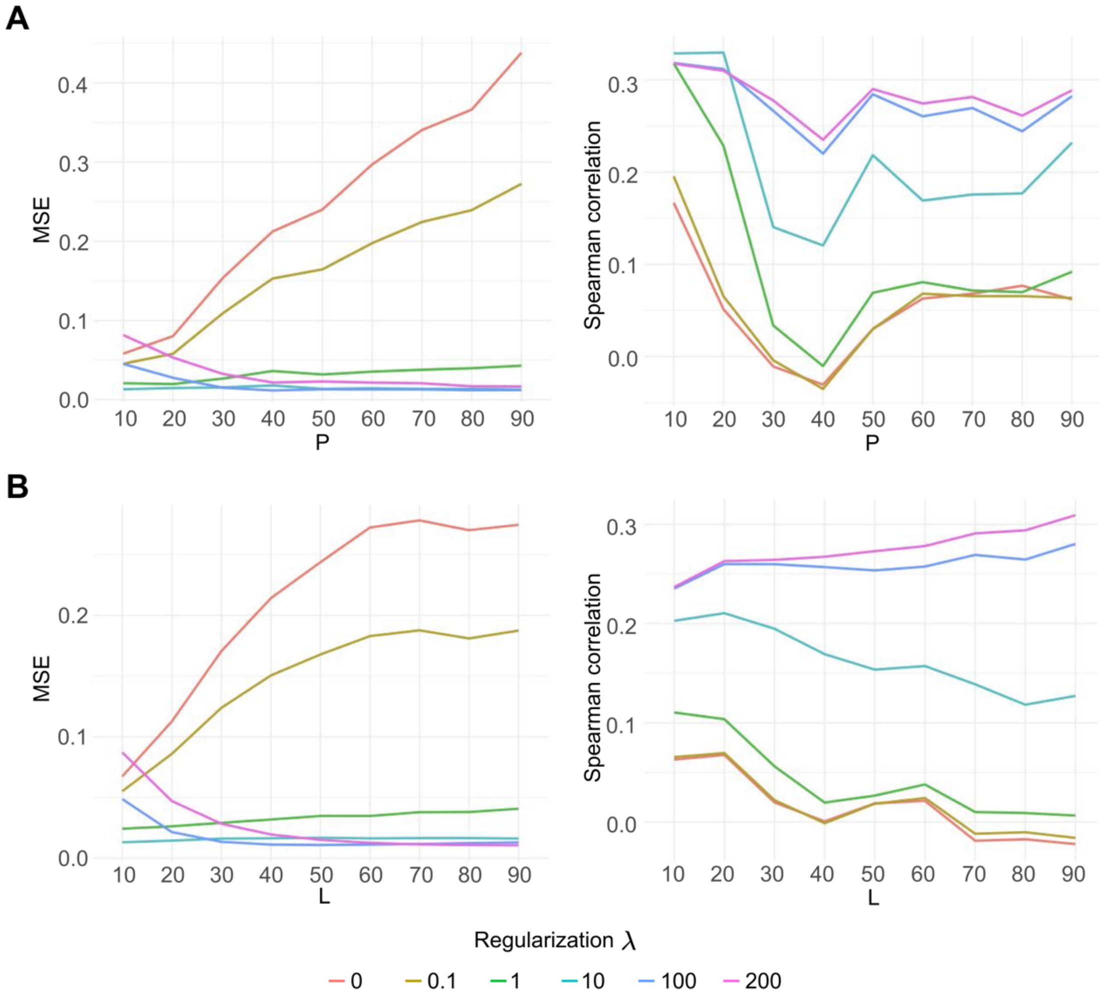


**Figure S11. L2 regularization of Granger causal model prevents overfitting**

A. Lineplots display the predicted mean-squared error (MSE, left) and Spearman correlation (right) on test data (see Results) as a function of the number of regulators. We used six different settings for the L2 regularization parameter.

B. Lineplots display the predicted mean-squared error (MSE, left) and Spearman correlation (right) on test data (see Results) as a function of the maximum lagged time stamps (*L*). We used six different settings on L2 regularization parameter.

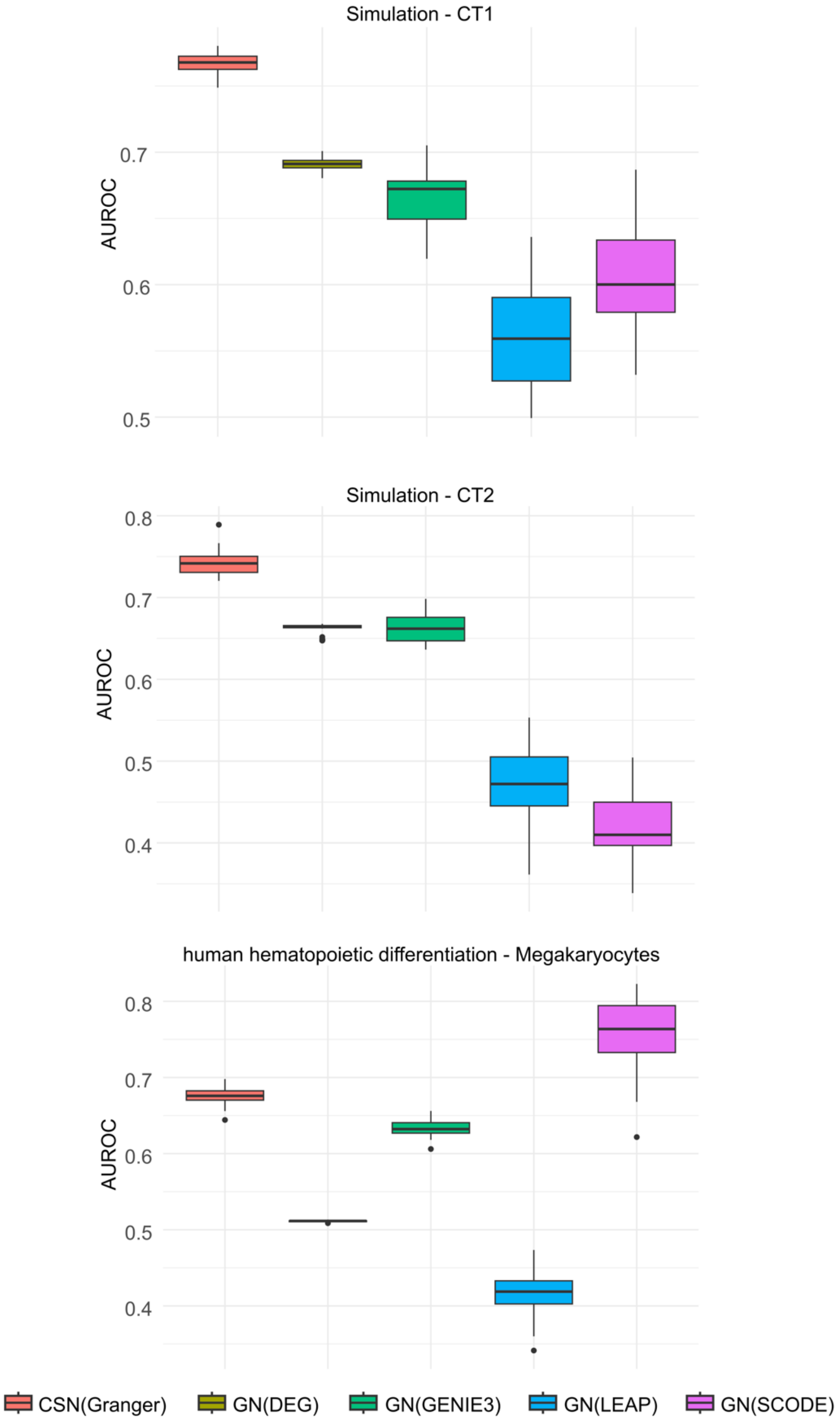


**Figure S12. Benchmarking lineage-specific GRN inference**

Boxplots showing the prediction performance evaluated by AUROC for five methods on lineage-specific GRN prediction.

The box in the boxplot represents the interquartile range (IQR).The whiskers extend to the smallest and largest values within 1.5 times the IQR. The black line within the box indicates the median.

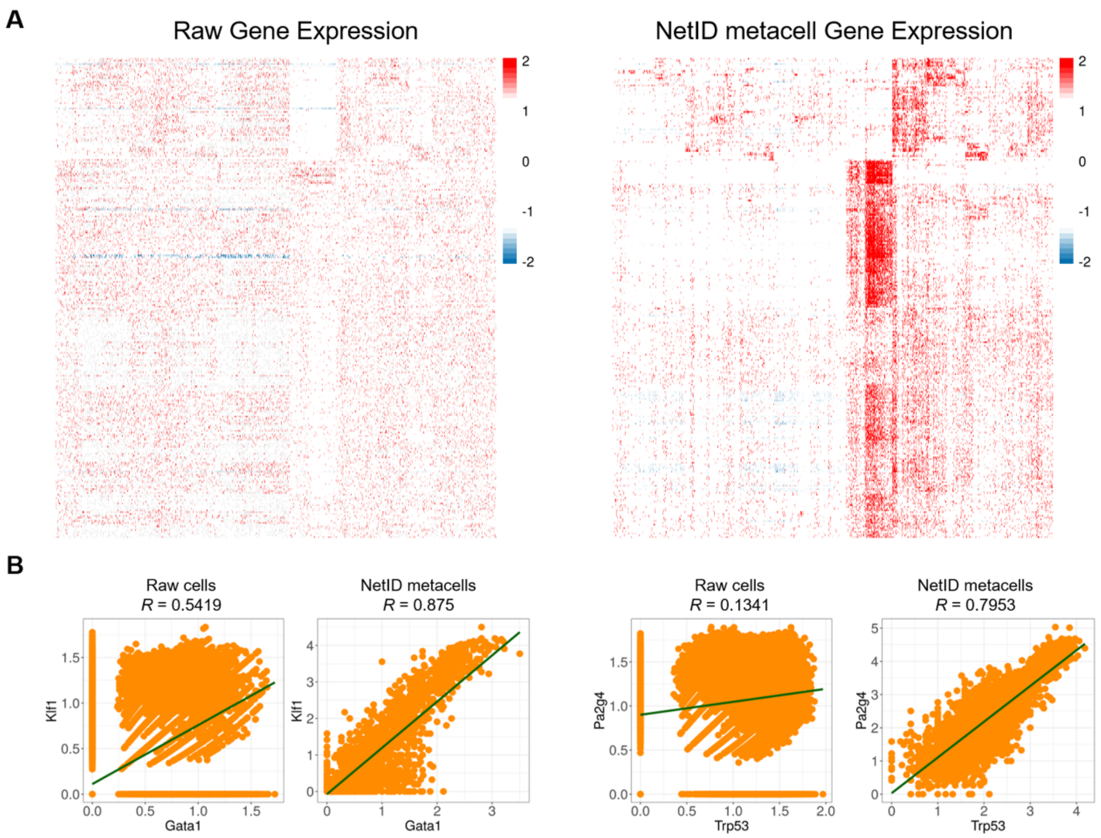


**Figure S13. Transcription Factor Expression Analysis for mouse hematopoiesis**

A. Heatmaps showing raw gene expression (left) and NetID inferred gene expression (right) for 271 transcription factors across sampled seed cells for the mouse hematopoiesis dataset [4].

B. Scatterplots showing the correlation between *Gata1* and *Klf1* (left) or between *Trp53* and *Pa2g4* (right) using raw gene expression or NetID metacell gene expression.

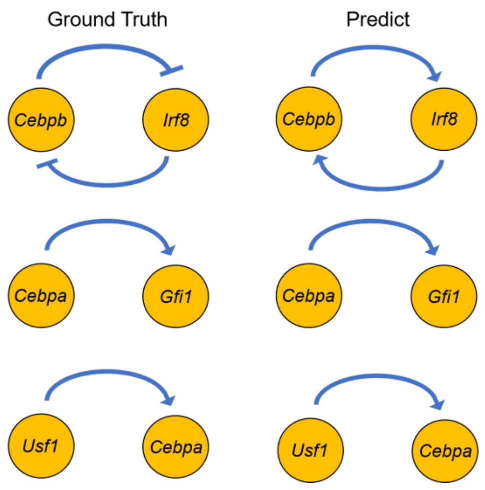


**Figure S14. Validation of NetID predicted gene regulatory relationships.**
NetID correctly predicted positive regulatory relationships from *Cebpa* to *Gfi1* [5], *Usf1* to *Cebpa* [6], *Xbp1* to *Cebpa* [7], and negative regulatory relationships from *Cebpa* to *Mycn* [8]. NetID incorrectly predicted a double negative feedback loop motif between *Cebpb* and *Irf8* [9] as double positive feedback loop motif.
